# *Shigella flexneri* undergoes niche-specific adaptation during infection

**DOI:** 10.64898/2026.09.15.751907

**Authors:** Yuang Sun, David W. Lazinski, Lauren K. Yum, Andrew Camilli, Hervé Agaisse

## Abstract

*Shigella flexneri* is a leading cause of diarrheal disease worldwide, yet bacterial factors required within distinct host niches remain poorly understood. Here, we used genome-wide transposon sequencing in an infant rabbit model of shigellosis to identify genes promoting bacterial fitness *in vivo*. The screen identified virulence factors on the large virulence plasmid and novel chromosomal fitness factors. Characterization of *zitB*, encoding a cation diffusion facilitator family Zn transporter, revealed a niche-specific fitness contribution. The Δ*zitB* mutant showed no growth defect *in vitro* in rich media or *in vivo* during the early infection phase (8 hpi) in epithelial cells. However, bacterial burden was reduced during the late infection phase (24 hpi) *in vivo*, when bacteria interact with immune cells, in both competitive and mono-infection assays. Reduced bacterial burden was associated with increased *MARCO*-specific macrophages, suggesting impaired colonization and killing by the Δ*zitB* mutant. Consistently, Δ*zitB* fitness was impaired in THP-1-derived macrophages. Metal chelation increased bacterial burden, indicating a role for macrophage-mediated metal toxicity. The Δ*zitB* mutant also showed increased sensitivity to zinc and copper. Finally, macrophage depletion *in vivo* restored bacterial burden to wild-type levels. Together, these findings identify ZitB as a niche-specific bacterial factor promoting *S. flexneri* fitness during macrophage-associated metal stress.

## Introduction

For decades, shigellosis, an infection of the intestinal tract caused by *Shigella spp.*, has remained a leading cause of diarrheal disease and mortality worldwide in children younger than 5 years^1,2^. Shigellosis disproportionately burdens low-income and mid-income countries where water quality and sanitation are poor are poor^3^, while in high-income countries, infection is typically associated with travel to high-risk regions and sporadic foodborne outbreaks^4^. *Shigella flexneri* and *Shigella sonnei* are the two most common etiologic agents of shigellosis^3^. *S. flexneri* has an extremely low infectious dose (∼100 cells) required for transmission^5^ via person-to-person contact. Antibiotics are the standard treatment for *S. flexneri* infection. In recent years, however, antimicrobial resistance has continued to rise, including the emergence of multidrug-resistant strains, underscoring the need for a better understanding of its pathogenesis^6–8^.

Foundational studies in non-human primates demonstrated that *S. flexneri* infection is marked by invasion of the intestinal epithelium, resulting in acute colitis in the mucosa and hemorrhage in the colon^9^. Subsequent seminal studies in tissue cultures defined a detailed framework for how *S. flexneri* invades epithelial cells and spreads from cell to cell^10–13^. Notably, the type III secretion system (T3SS), encoded by the large “invasion plasmid” (pINV)^14^, plays a critical role in pathogenesis, including manipulation of the host cellular processes to achieve actin-based motility and cell-to-cell spread^15,16^, modulation of signaling pathways such as inhibition of NF-κB activation, and suppression of innate immune responses including the induction of macrophage cell death^17,18^.

Despite numerous advances, after decades of efforts, there are still no effective small molecule drugs or licensed vaccines against *Shigella*. The lack of a reliable *in vivo* model was one of the major hindrances. Historically, animal models have been limited in their ability to recapitulate the full spectrum of shigellosis. Mice do not naturally develop diarrhea or ulcerative colitis^19,20^, and the Sereny test in the guinea pig’s eye assesses only localized inflammation at a non-physiological site^21^. Recently, work conducted in an *Nlrc4*^−/−^ mouse model revealed that disruption of the NAIP–NLRC4 inflammasome confers susceptibility^22^. Concurrently, infant rabbit models of shigellosis that are naturally susceptible to *S. flexneri* infection were developed^23,24^. These new models recapitulate key features of human shigellosis, including bloody diarrhea, mucosal ulceration, and immune cell infiltration, and thus provide systems in which the broader course of *S. flexneri* infection can be studied. Notably, in the infant rabbit model, shigellosis consist in a two-stage disease whereby bacteria first invade epithelial cells and spread from cell to cell. This early phase of infection (8 hpi) leads to epithelial fenestration and vascular lesions. During the late phase of infection (24 hpi), bacteria gain access to the lamina propria where they encounter immune cells, including neutrophils and macrophages^25^. *S. flexneri* must adapt to this challenging environment and the need for specific *S. flexneri* fitness factors might arise.

Genome-wide approaches such as transposon insertion sequencing (Tn-seq)^26,27^ allow systematic identification of bacterial genes that contribute to virulence and fitness under defined conditions^28–32^. To our knowledge, Tn-seq has not been applied to a model that recapitulates the full pathological features of shigellosis. Here, we applied high-density Tn-seq to an established infant rabbit model to identify bacterial factors required for *S. flexneri* infection *in vivo*. The screen identified canonical plasmid-encoded virulence genes associated with epithelial invasion, cell-to-cell spread, and immune evasion, while also revealing chromosomal genes with roles that could not be detected in epithelial cell assays. A mutant lacking one such chromosomal gene, *zitB*, encoding a Zn efflux transporter, showed no detectable phenotype in epithelial cell infection but was attenuated *in vivo*, prompting further investigation into its function in interactions with innate immune responses. These findings provide a foundation for understanding bacterial requirements at specific stages of shigellosis and highlight zinc efflux as a previously underappreciated survival strategy employed by *S. flexneri in vivo*.

## Results

### Tn-seq in the infant rabbit model of shigellosis identifies plasmid-encoded and chromosomal virulence and fitness factors

To identify bacterial factors required for shigellosis *in vivo*, we constructed a high-density transposon insertion library and performed Tn-seq using the infant rabbit model^23^. The library contained approximately 230,000 unique insertion sites, corresponding to an average density of approximately one insertion every 21 bp. Output populations recovered from four independently infected rabbits were compared with the input library. Analysis of the input library predicted 546 essential chromosomal genes and no essential genes on the invasion plasmid (pINV) (Supplementary Data 1), largely consistent with a previous Tn-seq analysis of *S. flexneri* 2457T that identified 481 ORFs required for robust growth in rich medium^30^. Complete processed input and output values and statistical results are provided in Supplementary Data 2.

The screen identified 47 pINV-encoded genes for which transposon insertions were significantly depleted in the output relative to the input library (Fig. 1a and Supplementary Data 1). These include *spa* and *mxi* structural genes encoding the type III secretion system (T3SS) within the pINV entry region^14,33^ as well as genes located outside this region, including virulence regulators *virF* and *virB*; and *icsA* required for cell-to-cell spread^12,34^. Of note, 33 of the 37 genes within the entry region were confirmed hits in the Tn-seq, highlighting the importance of this pathogenicity island (Fig. 1c). Because two entry region genes *orf131a* and *orf131b*^35^ were not annotated as features in the pINV reference annotation used for the automated workflow, these two open reading frames (ORFs) were analyzed separately and included manually in the entry-region analysis. Although found within the entry region defined by low GC-content, *orf131a* and *orf131b* located on the high-coordinate end, together with *ipaJ* and *acp* on the low-coordinate end were not identified as significant hits (Supplementary Fig. 1 and Supplementary Data 2).

**Fig. 1:**
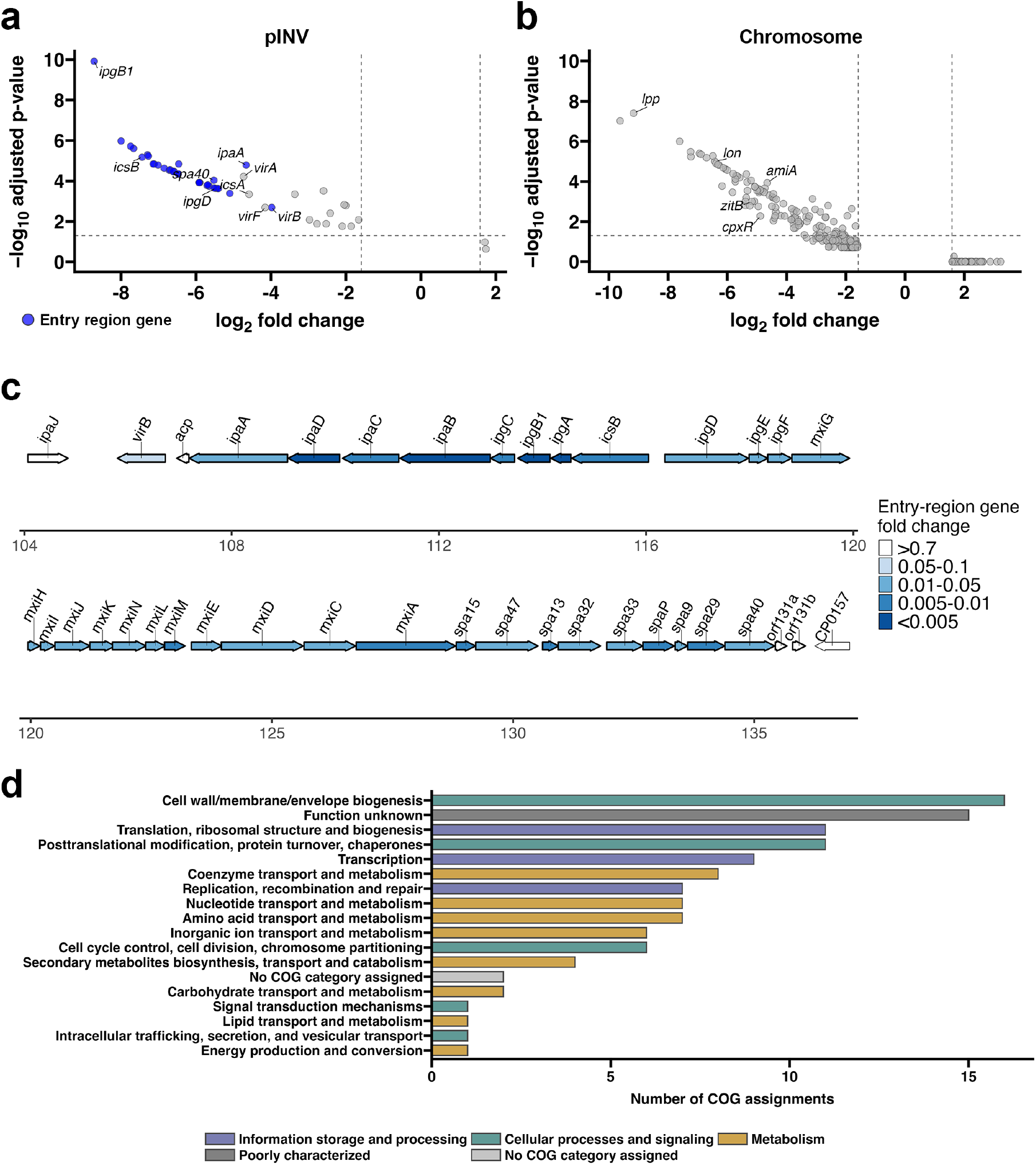
Tn-seq analysis identifies *S. flexneri* virulence and fitness factors encoded on pINV and the chromosome. **a**–**b** Volcano plots showing the relationship between log_2_ mean output/input ratio (fold change) and −log_10_ Holm-adjusted *P* value for pINV genes (**a**) and chromosomal genes (**b**) identified in the Tn-seq screen. For each gene, output/input ratios were calculated for four independently infected rabbits (*n* = 4). Selected genes are labeled. In **a**, genes within the pINV entry region are highlighted in blue. Genes with mean output/input ratios below 1/3 or above 3 were tested against a null ratio of 1 using one-sample, one-tailed *t*-tests, followed by Holm correction for multiple comparisons. Vertical dashed lines indicate ratio cutoffs of 1/3 and 3, and the horizontal dashed line indicates an adjusted *P* value of 0.05. **c** Gene map of the pINV entry region on plasmid NC_004851.1, with genomic coordinates shown. Genes are colored according to the output/input ratio (fold change) categories indicated. **d** COG functional-category assignments of significant chromosomal hits, colored by broad functional group. Genes assigned to multiple COG categories were counted in each applicable category.

In addition to pINV-encoded genes, the screen identified 110 chromosomal genes that showed statistically significant reductions in output abundance relative to the input (Fig. 1b and Supplementary Data 1). Clusters of Orthologous Genes (COG) annotation designated these genes across diverse cellular functions, with the largest categories comprising cell wall, membrane and envelope biogenesis; proteins of unknown function; translation and ribosome structure and biogenesis; and post-translational modification, protein turnover and chaperones (Fig. 1d). To examine functional associations among the chromosomal hits, we analyzed the corresponding proteins using the Search Tool for the Retrieval of Interacting Genes/Proteins (STRING). STRING recognized 109 of the 110 submitted proteins. After disconnected proteins were omitted, 59 proteins formed a network of 64 functional associations. The Markov cluster algorithm (MCL) grouped the 59 proteins into 20 non-overlapping clusters based on network connectivity. These included clusters associated with lipopolysaccharide biosynthesis (*waaI*, *waaY*, *waaQ*, *waaP*, *waaL*, *waaJ*, and *waaD*), outer-membrane lipid homeostasis (*vacJ*, *vpsA*, *vpsB*, *vpsC*, and *yrbC*), sulfur metabolism (*cysC*, *cysJ*, *cysQ*, and *cysH*), stress responses and protein quality control (*dnaJ*, *lon*, *hslU*, and *hslV*), homologous recombination (*ybaB*, *recR*, *ruvA*, and *recF*), and cell division (*minC*, *minD*, and *minE*) (Supplementary Fig. 2 and Supplementary Data 3).

### Disruption of selected Tn-seq candidate genes differentially affects epithelial cell-to-cell spread

Cell-to-cell spread is a central step in *S. flexneri* pathogenesis and is known to depend on pINV-encoded genes^34,36,37^, many of which were also identified as significant hits in our Tn-seq screen, such as *ipgB1*, *ipgD*, and *icsA* (Fig. 1a). To explore whether chromosomal genes could likewise influence this process, we examined bacterial dissemination in HT-29 cell monolayers using selected chromosomal candidates representing diverse functions: *cpxR*, a regulator of *virF* expression^38^; *amiA* and *waaY*, involved in peptidoglycan remodeling and lipopolysaccharide core phosphorylation, respectively^39,40^; *zitB*, encoding a zinc exporter^41^; *hslV*, encoding the protease component of the HslUV complex^42^; and *hha*, encoding an H-NS-interacting transcriptional modulator^43^. All six genes had mean output/input ratios below 0.33 (i.e. log_2_ fold change < −1.585); *cpxR*, *amiA*, *zitB*, *hslV*, and *waaY* were significantly depleted after correction for multiple comparisons, whereas *hha* did not meet the significance threshold (Holm-adjusted *P* = 0.067). The mutants for all six genes showed similar growth kinetics to WT in lysogeny broth (LB), although Δ*cpxR* and Δ*hha* showed modest but statistically significant reductions in area under the curve (AUC) of growth curves (Supplementary Fig. 4a–b).

In HT-29 cells, Δ*cpxR*, as expected of a *virF* regulator, produced no detectable foci (Fig. 2a, c and Supplementary Fig. 3b). Δ*amiA* formed significantly smaller foci than WT, with a mean focus area of 3371 px^2^ versus 4187 px^2^ for WT, corresponding to an approximately 20% reduction (Fig. 2a, c). Elongated bacterial structures were observed within Δ*amiA* foci, suggestive of cell chaining (Fig. 2b). Δ*hha* also showed a significant but more modest reduction in focus area (Fig. 2c and Supplementary Fig. 3a–b). By contrast, Δ*zitB*, Δ*hslV*, and Δ*waaY* showed no detectable differences in focus area compared to WT (Fig. 2a, c and Supplementary Fig. 3a–b). Thus, disruption of the selected chromosomal candidates resulted in phenotypes ranging from complete loss of focus formation in Δ*cpxR*, to partial defects in Δ*amiA* and Δ*hha*, to no detectable phenotype in Δ*zitB*, Δ*hslV*, and Δ*waaY*.

**Fig. 2:**
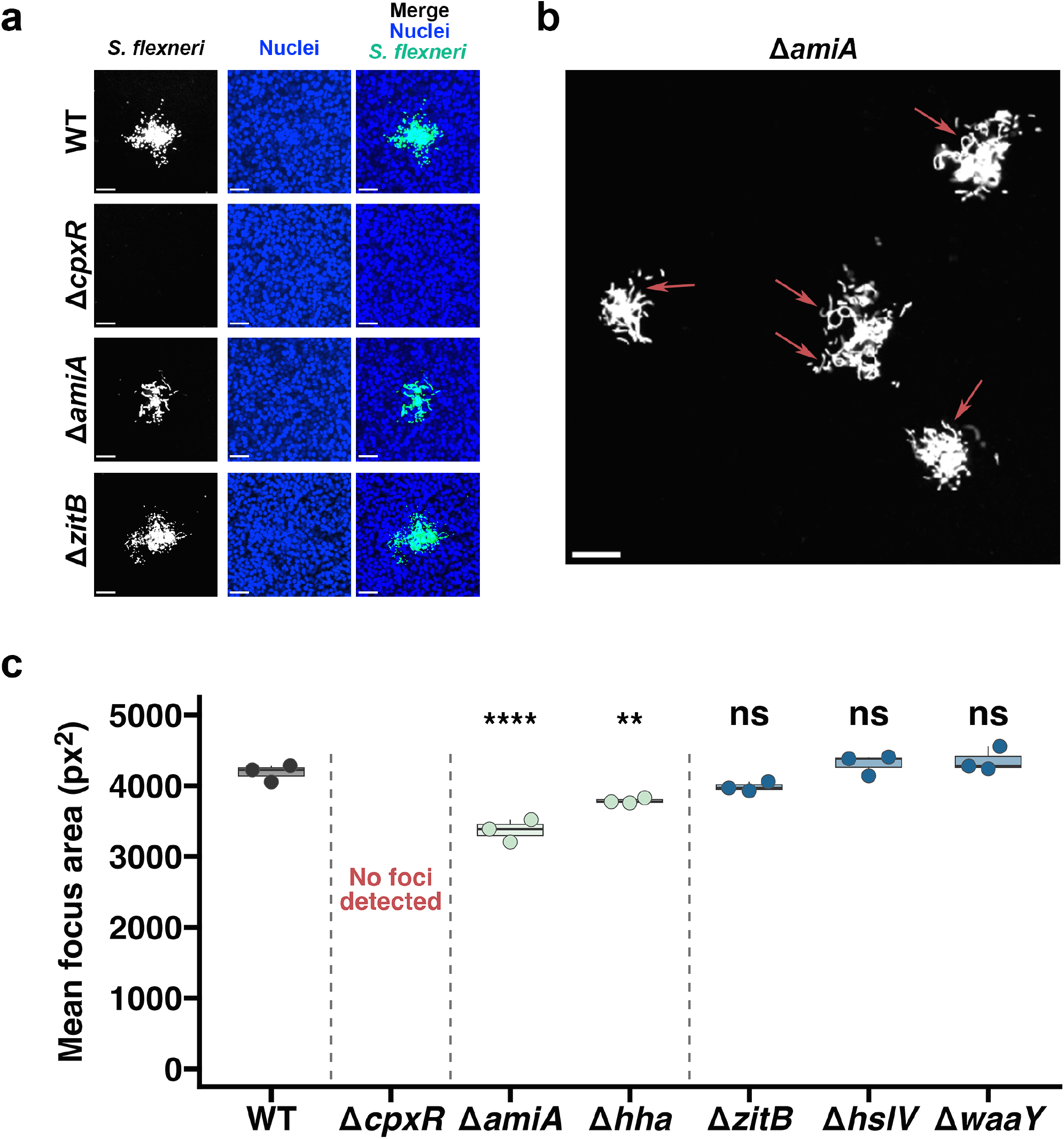
Assessment of selected Tn-seq candidate mutants during HT-29 cell-to-cell spread. **a** Representative images of infection foci formed by WT and the indicated mutants in HT-29 cell monolayers at 8 hpi. *S. flexneri* is shown in white in the single-channel images and green in the merged images; nuclei are shown in blue. **b** Additional representative image of Δ*amiA* infection foci. Arrows indicate elongated bacterial structures suggestive of cell chaining. **a**–**b** Scale bars, 50 μm. **c** Quantification of mean focus area for WT and the indicated mutants. Each dot represents one independent experiment (*n* = 3). Boxes show the median and interquartile range (IQR); whiskers extend to the smallest and largest observations within 1.5 × IQR of the lower and upper quartiles, respectively. Δ*cpxR* produced no detectable foci. Statistical analysis was performed using linear regression followed by Dunnett-adjusted comparisons with WT. \*\**P* < 0.01; \*\*\*\**P* < 0.0001; ns, not significant.

### *zitB* is a fitness factor for sustained bacterial burden *in vivo*

Although Δ*zitB*, Δ*hslV*, and Δ*waaY* were identified as hits in the screen, none showed a detectable defect in cell-to-cell spread in HT-29 cells, suggesting that they may contribute to infection through mechanisms independent of dissemination. Given the role of ZitB in zinc efflux^44,45^, we prioritized *zitB* for further study as a candidate fitness factor that may be specifically required *in vivo*. To validate the competitive defect identified by Tn-seq using an independent readout, we performed 1:1 competitive infection with WT and Δ*zitB* in infant rabbits. WT and Δ*zitB* showed comparable growth in tryptic soy broth (TSB), the medium used for infection inoculum preparation, with no significant difference in growth as assessed by AUC (Supplementary Fig. 4c–d). At 24 hpi, Δ*zitB* was outcompeted by WT, with a competitive index of 0.156, corresponding to an approximately 6.4-fold competitive disadvantage relative to WT (Fig. 3a).

**Fig. 3:**
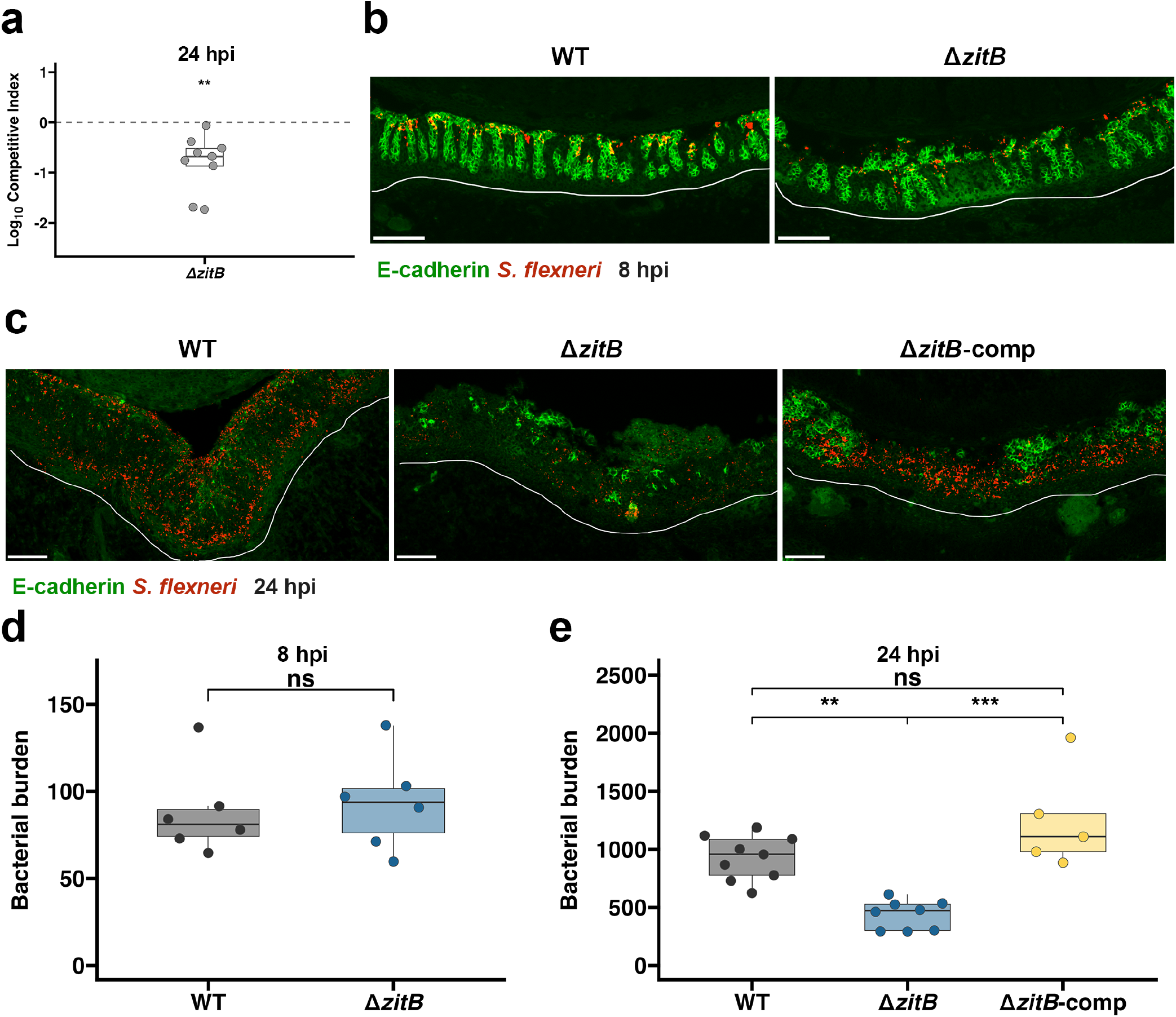
ZitB promotes sustained *S. flexneri* burden during infant rabbit infection. **a** Log_10_ competitive index of Δ*zitB* relative to WT at 24 hpi following 1:1 mixed infection. Each dot represents one rabbit (*n* = 9); the dashed line indicates a neutral competitive index. The mean log_10_ competitive index was compared with 0 using a linear model. **b** Representative images of colonic tissue from rabbits infected with WT or Δ*zitB* at 8 hpi. **c** Representative images of colonic tissue from rabbits infected with WT, Δ*zitB*, or Δ*zitB*-comp at 24 hpi. In **b**–**c**, *S. flexneri* is shown in red, E-cadherin in green, and white lines delineate the boundary between the mucosa and submucosa. Scale bars, 100 μm. **d** Quantification of bacterial burden at 8 hpi. Each dot represents the model-predicted mean for one rabbit (*n* = 6 per strain). **e** Quantification of bacterial burden at 24 hpi. Each dot represents the model-predicted mean for one rabbit (WT, *n* = 9; Δ*zitB*, *n* = 8; Δ*zitB*-comp, *n* = 5). For **d**–**e**, data were analyzed using linear mixed-effects models with strain as a fixed effect and rabbit as a random intercept. Pairwise comparisons in **e** were Tukey-adjusted. In **a, d, and e**, boxes show the median and IQR; whiskers extend to the smallest and largest observations within 1.5 × IQR of the lower and upper quartiles, respectively. \*\**P* < 0.01; \*\*\**P* < 0.001; ns, not significant.

Because competitive index does not necessarily match bacterial burden difference during mono-infection, we next examined WT and Δ*zitB* in mono-infected infant rabbits. To determine when the Δ*zitB* defect emerged, colon tissues were analyzed at 8 and 24 hpi by *S. flexneri* immunostaining and imaging-based quantification of tissue-associated bacterial burden (Supplementary Fig. 5). At 8 hpi, bacterial signal in tissues infected with WT or Δ*zitB* was largely associated with E-cadherin-positive epithelial cells near the luminal surface of the mucosa (Fig. 3b). Δ*zitB* exhibited bacterial burden comparable to that of WT (Fig. 3d), consistent with its preserved epithelial dissemination phenotype in HT-29 cells (Fig. 2a, c). By 24 hpi, bacterial signal was more broadly distributed throughout the mucosa, including within E-cadherin-negative regions (Fig. 3c); Δ*zitB* exhibited decreased bacterial burden compared with WT, with estimated marginal means of 437 and 927, respectively (Fig. 3e). To determine whether this defect resulted from the loss of *zitB*, we complemented Δ*zitB in trans* with *zitB* expressed under its native promoter, hereafter referred to as Δ*zitB*-comp. Complementation restored bacterial burden to levels comparable to WT: the estimated marginal mean for Δ*zitB*-comp was 1248, significantly higher than that of Δ*zitB* and not significantly different from WT (Fig. 3d, e).

Despite its lower bacterial burden at 24 hpi, infection with Δ*zitB* did not show detectable attenuation of overall pathology. Infant rabbits infected with WT or Δ*zitB* showed similar diarrhea and blood scores at both 8 and 24 hpi (Supplementary Fig. 6a, b, e, f). Histopathological analysis revealed progressive tissue pathology in both WT and Δ*zitB* infected rabbits: At 8 hpi, infection by both strains produced sporadic epithelial fenestration and focal hemorrhagic lesions within the mucosa. By 24 hpi, epithelial fenestration was widespread in both WT and Δ*zitB* infected tissues, accompanied by more prominent hemorrhage (Supplementary Fig. 6c, d, g, h). Quantification of epithelial fenestration showed no significant difference between the two strains at 24 hpi (Supplementary Fig. 6i). The preservation of epithelial fenestration in Δ*zitB*-infected tissues is consistent with its efficient epithelial dissemination in HT-29 cells (Fig. 2c) and bacterial burden comparable to WT at 8 hpi (Fig. 3d). Together, these findings indicate that *zitB* is not required during the initial epithelial phase of infection but is required to sustain bacterial burden during later infection.

### Δ*zitB* infection is associated with higher *MARCO* signal *in vivo*

The delayed emergence of the Δ*zitB* fitness defect suggested that ZitB might support bacterial fitness during interactions with host immune cells. To investigate the immune responses in step with the emergence of the Δ*zitB* burden defect, we examined neutrophil and macrophage responses at both 8 and 24 hpi using RNA *in situ* hybridization (RNAscope) and the imaging-analysis workflow described in Supplementary Fig. 5.

Neutrophils are major innate immune cells recruited during *Shigella* infection^46^. To assess neutrophil responses, we quantified *LCN2*, which is tightly associated with neutrophils during acute infection^25,47^. At 8 hpi, when bacterial burdens were comparable between WT and Δ*zitB*, normalized mucosal *LCN2* signal did not differ between the two groups (Supplementary Fig. 7a). At 24 hpi, both WT and Δ*zitB* infections elicited robust *LCN2* signal. In both strains, the mucosa accounted for more than 80% of the estimated compartmental distribution of *LCN2*-positive signal, with approximately 10% in the submucosa and less than 5% in the muscle (Fig. 4c). Normalized *LCN2* signal in the mucosa, submucosa and muscle did not differ significantly between rabbits infected with WT or Δ*zitB* (Fig. 4a–b and Supplementary Fig. 7c–d).

**Fig. 4:**
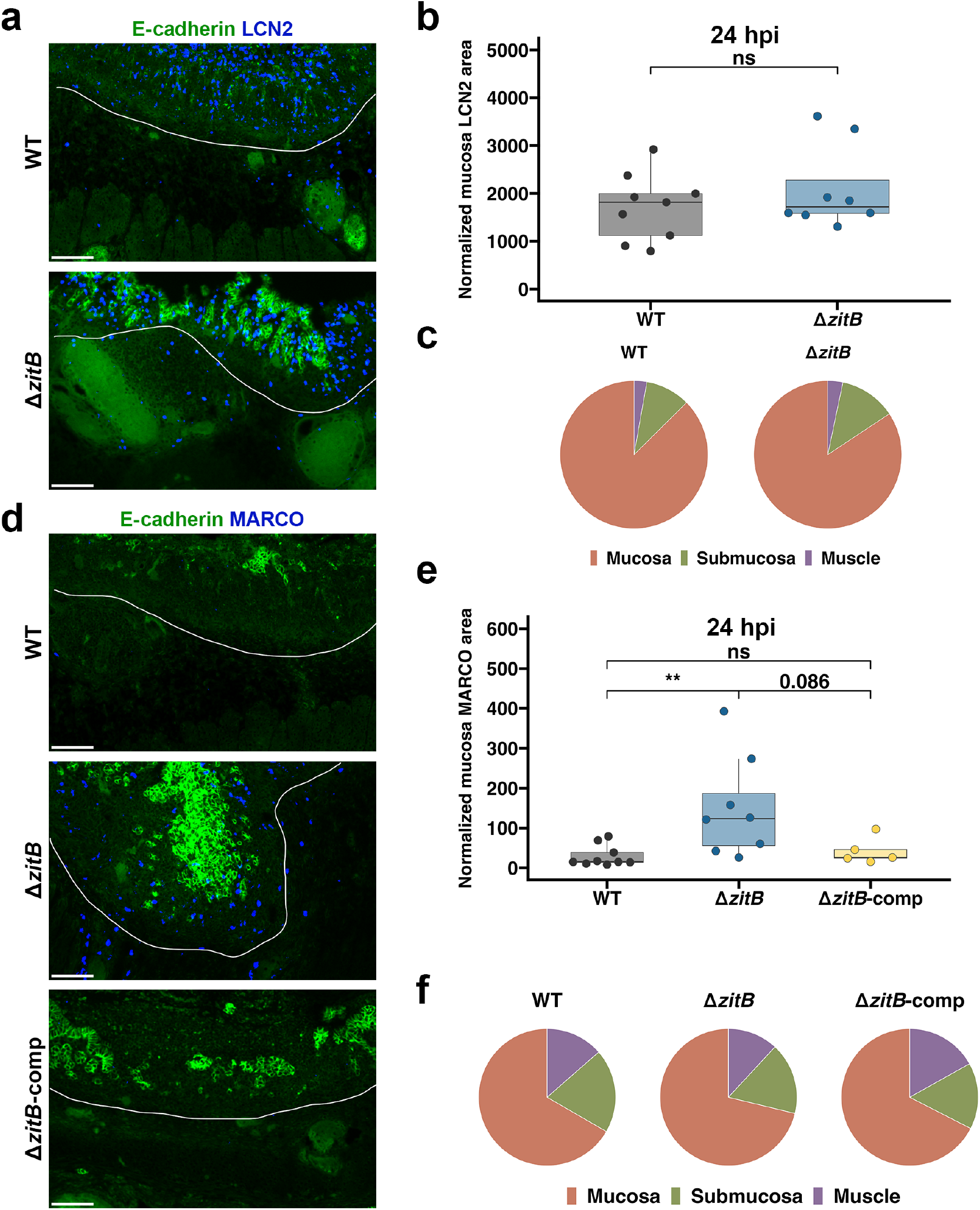
WT and Δ*zitB* infections exhibit differential mucosal *MARCO* signal *in vivo*. **a** Representative images of colonic tissue from rabbits infected with WT or Δ*zitB* at 24 hpi showing *LCN2* and E-cadherin. **b** Quantification of normalized mucosal *LCN2*-positive area. Each dot represents the model-predicted mean for one rabbit (WT, *n* = 9; Δ*zitB*, *n* = 8). **c** Estimated compartmental distribution of *LCN2*-positive area among the mucosa, submucosa, and muscle. **d** Representative images of colonic tissue from rabbits infected with WT, Δ*zitB*, or Δ*zitB*-comp at 24 hpi showing *MARCO* and E-cadherin. **e** Quantification of normalized mucosal *MARCO*-positive area. Each dot represents the model-predicted mean for one rabbit (WT, *n* = 9; Δ*zitB*, *n* = 8; Δ*zitB*-comp, *n* = 5). **f** Estimated compartmental distribution of *MARCO*-positive area among the mucosa, submucosa, and muscle. In **a** and **d**, *LCN2* or *MARCO* is shown in blue, E-cadherin in green, and white lines delineate the boundary between the mucosa and submucosa. Scale bars, 100 μm. For **b** and **e**, data were analyzed using linear mixed-effects models with strain as a fixed effect and rabbit as a random intercept. Pairwise comparisons in **e** were Tukey-adjusted. Boxes show the median and IQR; whiskers extend to the smallest and largest observations within 1.5 × IQR of the lower and upper quartiles, respectively. \*\**P* < 0.01; ns, not significant.

We next examined macrophage-associated responses in infected colons. Previous work showed that MARCO is a specific representative marker of the macrophage population^25^. MARCO is a macrophage-expressed class A scavenger receptor that contributes to pathogen recognition and opsonin-independent phagocytosis of bacteria^48,49^. At 8 hpi, normalized mucosal *MARCO* did not differ between WT and Δ*zitB* infections. At 24 hpi, however, normalized mucosal *MARCO* signal was higher in tissues infected with Δ*zitB* than in those infected with WT, with estimated marginal means of 150.1 and 29.8, respectively.

Complementation of Δ*zitB* reduced *MARCO* signal toward WT levels, with Δ*zitB*-comp exhibiting an estimated marginal mean of 41.8. While the comparison between Δ*zitB* and Δ*zitB*-comp did not reach statistical significance (Tukey-adjusted *P* = 0.086), Δ*zitB*-comp was not significantly different from WT (Fig. 4d–e). Normalized *MARCO* signal was also higher during Δ*zitB* infection in the submucosa and muscle compared to WT, whereas Δ*zitB*-comp did not differ significantly from WT or Δ*zitB* in either compartment (Supplementary Fig. 7e–f). In all three strains, *MARCO* signal was localized primarily to the mucosa, which accounted for approximately 70% of its estimated compartmental distribution; the submucosa and muscle accounted for more than 15% and 10%, respectively (Fig. 4f). Together, these data show that *LCN2*-and *MARCO*-associated signals were concentrated primarily in the mucosa. While WT and Δ*zitB* elicited comparable *LCN2* responses, Δ*zitB* infection was associated with higher mucosal *MARCO* signal, suggesting a differential interaction with macrophages.

### ZitB supports survival in macrophages and resistance to zinc and copper stress

Next, we asked whether loss of *zitB* altered the direct interaction of *S. flexneri* with macrophages. THP-1-derived macrophages were infected with WT, Δ*zitB* or Δ*zitB*-comp, and bacterial CFU were quantified at 2 and 4 hpi. Fewer CFU were recovered from Δ*zitB* than from WT at both time points (Supplementary Fig. 8g). Because CFU recovery varied among experimental batches, values were normalized to WT within the corresponding batch. Following normalization, Δ*zitB* CFU recovery was significantly lower than WT at both 2 and 4 hpi, whereas Δ*zitB*-comp did not differ significantly from WT (Fig. 5a). These findings indicate that ZitB may promote bacterial survival during macrophage infection.

**Fig. 5:**
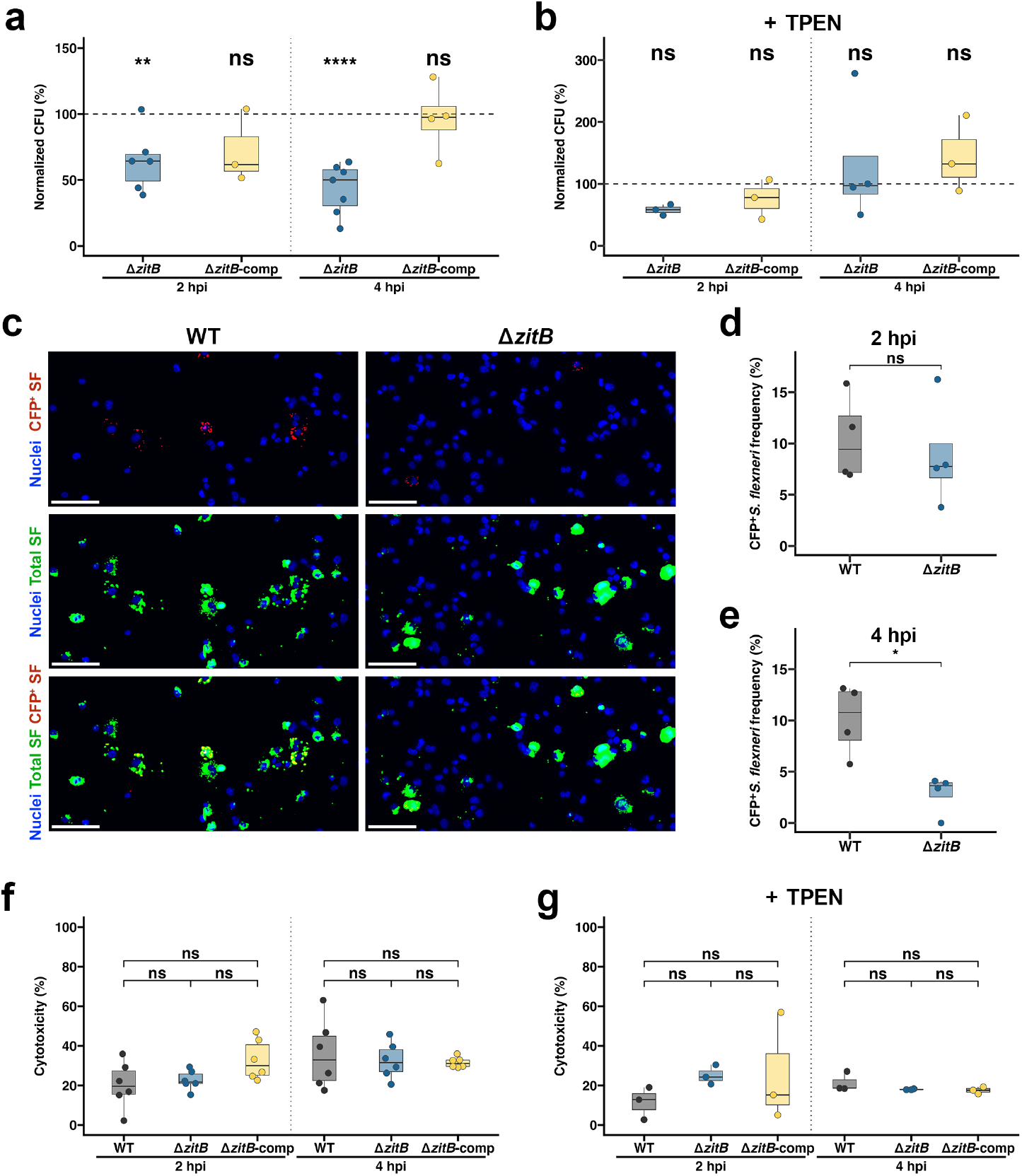
ZitB promotes *S. flexneri* survival in THP-1-derived macrophages. **a** THP-1-derived macrophages were infected with WT, Δ*zitB*, or Δ*zitB*-comp, and CFU were recovered at 2 and 4 hpi. Δ*zitB* and Δ*zitB*-comp values were expressed as percentages of the batch-matched WT value. Each dot represents one independent infection. At 2 and 4 hpi, respectively, *n* = 8 and 9 for WT, 6 and 7 for Δ*zitB*, and 3 and 4 for Δ*zitB*-comp. The dashed line indicates the WT reference value of 100%. **b** The assay in **a** was repeated with 10 μM TPEN added at the start of infection; CFU were recovered and normalized to batch-matched WT as in **a**. At 2 and 4 hpi, respectively, *n* = 3 and 4 for WT, 3 and 4 for Δ*zitB*, and 3 and 3 for Δ*zitB*-comp. **c** Representative images of macrophages infected with WT or Δ*zitB* at 4 hpi. SF denotes *S. flexneri*. Nuclei are shown in blue, total SF detected by anti-*Shigella* immunostaining in green, and CFP^+^ SF in red. Scale bars, 100 μm. **d**–**e** Frequency of CFP-expressing (CFP^+^) *S. flexneri* detection at 2 hpi (**d**) and 4 hpi (**e**). Each dot represents one independent infection (*n* = 4 per strain). **f**–**g** Cytotoxicity following infection with WT, Δ*zitB*, or Δ*zitB*-comp without (**f**) or with 10 μM TPEN (**g**), measured by LDH release at 2 and 4 hpi. Each dot represents one independent infection (*n* = 6 in **f** and *n* = 3 in **g** per strain at each time point). Cytotoxicity was calculated relative to spontaneous and maximum-release controls. For **a**–**b** and **f**–**g**, data were analyzed separately by time point and TPEN condition using linear models followed by Tukey-adjusted pairwise comparisons of estimated marginal means; only comparisons with WT are shown in **a**–**b**. For **d**–**e**, strains were compared separately at each time point using linear models. For **a**–**b** and **d**–**g**, boxes show the median and IQR; whiskers extend to the smallest and largest observations within 1.5 × IQR of the lower and upper quartiles, respectively. \**P* < 0.05; \*\**P* < 0.01; \*\*\*\**P* < 0.0001; ns, not significant.

Because ZitB is a predicted zinc exporter and disruption of bacterial zinc efflux can increase susceptibility to zinc intoxication^44,45,50,51^, we next examined the growth of WT, Δ*zitB*, and Δ*zitB*-comp under defined metal conditions. All three strains reached comparable endpoint CFU in M9 medium with no added metal (Supplementary Fig. 8a). Addition of 25 μM Zn^2+^ reduced Δ*zitB* CFU compared to WT, and complementation restored CFU to levels comparable to WT (Supplementary Fig. 8b). This difference was no longer detected when the chelator N,N,N′,N′-Tetrakis(2-pyridylmethyl)ethylenediamine (TPEN) was added together with Zn^2+^ in equimolar amounts (Supplementary Fig. 8c). Because cation diffusion facilitator (CDF)-family transporters may exhibit broad specificity for divalent metals^52^, we also examined susceptibility to other transition metals. Δ*zitB* growth was also impaired in M9 supplemented with 25 μM Cu^2+^, and this phenotype was also restored by complementation (Supplementary Fig. 8d). By contrast, WT and Δ*zitB* showed comparable growth following supplementation with Fe^2+^ or Mn^2+^ (Supplementary Fig. 8e–f). Thus, loss of *zitB* increased susceptibility to zinc and copper stress under minimal medium conditions.

Because macrophages can deploy transition metals, including zinc and copper, as antimicrobial agents^53,54^, we then tested whether metal chelation altered the Δ*zitB* phenotype in THP-1 macrophages. In the presence of TPEN, the normalized CFU recovered for Δ*zitB* and Δ*zitB*-comp did not differ significantly from WT at either 2 or 4 hpi (Fig. 5b and Supplementary Fig. 8h), suggesting that TPEN-chelatable transition metals, including zinc and copper, contribute to the Δ*zitB* phenotype in THP-1-derived macrophages.

Because CFP fluorescence delineated intact rod-shaped bacteria and was used as an indicator of active bacteria in previous studies^37,55^, we used CFP positivity as a proxy for viable bacteria, and anti-*S. flexneri* immunostaining for the total *Shigella* population to measure an imaging-based readout complementary to CFU recovery. At 2 hpi, the frequency of infected macrophages containing CFP-positive viable bacteria was comparable between WT and Δ*zitB*. At 4 hpi, this frequency was lower for Δ*zitB* than for WT (Fig. 5c–e). The reduced occurrence of viable Δ*zitB* was consistent with reduced fitness of Δ*zitB* during macrophage infection.

Finally, we tested whether reduced Δ*zitB* recovery was accompanied by altered macrophage cytotoxicity. Lactate dehydrogenase (LDH) release did not differ significantly among macrophages infected with WT, Δ*zitB*, or Δ*zitB*-comp at either 2 or 4 hpi, with or without TPEN treatment (Fig. 5f–g). Together, these findings indicate that ZitB supports *S. flexneri* survival within THP-1-derived macrophages and implicate chelatable transition metals in macrophage-associated Δ*zitB* fitness defect.

### Macrophage depletion abrogates the Δ*zitB* bacterial burden defect *in vivo*

We next asked whether macrophages contribute to the impaired Δ*zitB* fitness *in vivo*. Given the reduced Δ*zitB* survival within THP-1-derived macrophages and the inversely related bacterial burden and mucosal *MARCO* differences between WT and Δ*zitB* in infant rabbits, we hypothesized that macrophage depletion would rescue the *zitB* fitness defect and abrogate the difference in bacterial burden *in vivo*. To test this hypothesis, infant rabbits were treated with clodronate liposomes prior to infection, which are taken up by phagocytic cells and are widely used for depleting macrophages *in vivo*^56^. Rabbits treated with control liposomes were analyzed in parallel. In the control group, as previously shown in the absence of liposomes, bacterial burden was decreased in animals infected with the Δ*zitB* mutant at 24 hpi (Fig. 6a–b). This bacterial burden difference was abrogated in the clodronate group (Fig. 6a–b). The higher mucosal *MARCO* signal in Δ*zitB* infection was likewise preserved in the control group (Fig. 6a, c). Following clodronate liposome treatment, mucosal *MARCO* signal was low in both groups and did not differ between WT and Δ*zitB*, with estimated marginal means of 5.6 and 6.9, respectively (Fig. 6a, c). Mucosal *LCN2* signal remained robust after clodronate liposome treatment and did not differ significantly between WT and Δ*zitB* infections (Supplementary Fig. 9), indicating that the treatment did not diminish the *LCN2*-associated neutrophil response. Together, these findings support the notion that *MARCO*-expressing macrophages are necessary to mediate the observed defect in Δ*zitB* fitness *in vivo* at 24 hpi.

**Fig. 6:**
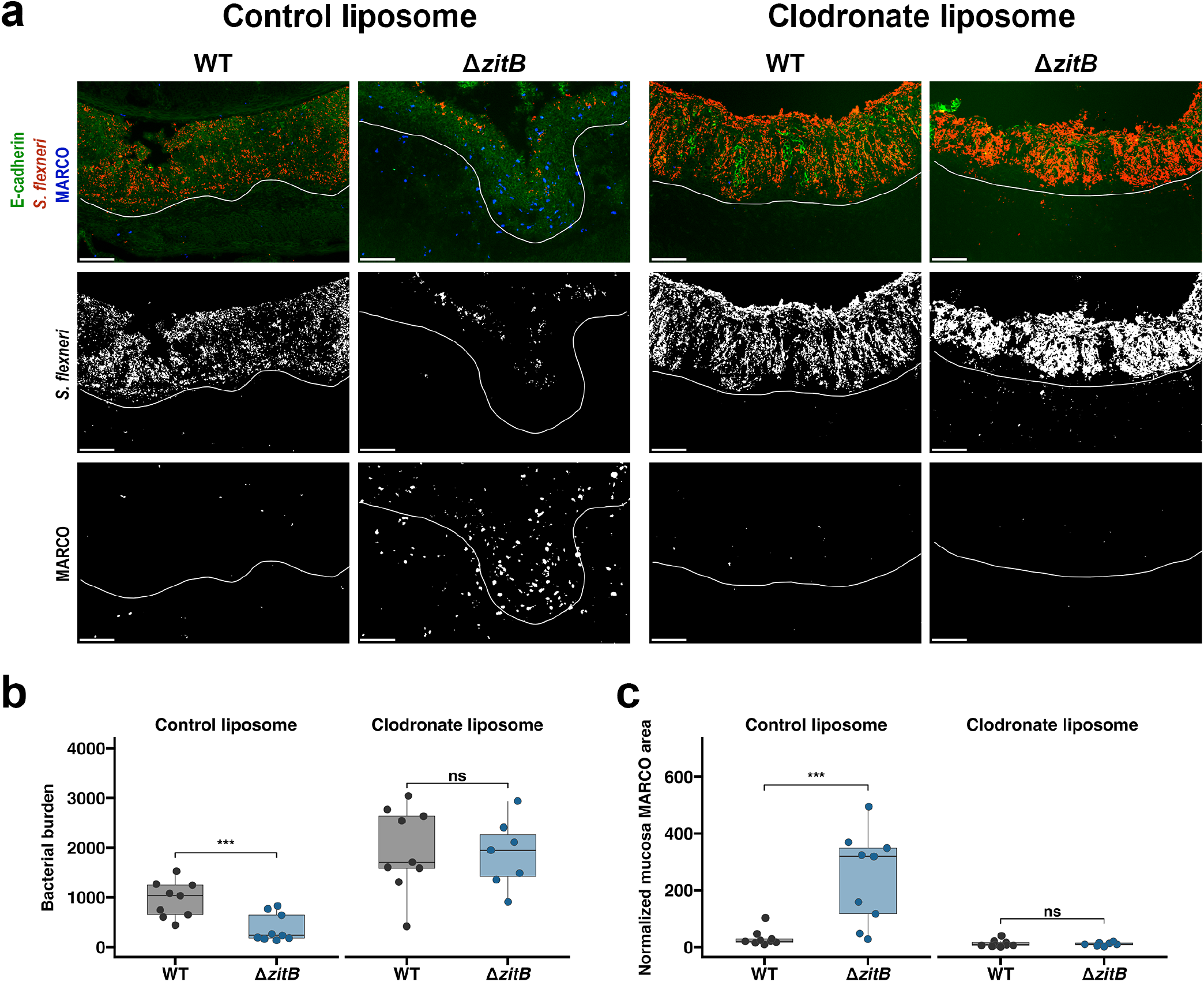
Clodronate liposome treatment abrogates WT–Δ*zitB* differences in bacterial burden and mucosal *MARCO* signal. **a** Representative merged and single-channel images of colonic tissue from rabbits treated intravenously with control or clodronate liposomes 48 h before infection with WT or Δ*zitB* and collected at 24 hpi. *S. flexneri* is shown in red, *MARCO* in blue, and E-cadherin in green. White lines delineate the boundary between the mucosa and submucosa. **b** Quantification of tissue-associated bacterial burden following control or clodronate liposome treatment. **c** Quantification of normalized mucosal *MARCO*-positive area following control or clodronate liposome treatment. For **b**–**c**, each dot represents the model-predicted mean for one rabbit on the untransformed scale. Sample sizes were *n* = 9 per strain following control liposome treatment and *n* = 9 for WT and *n* = 7 for Δ*zitB* following clodronate liposome treatment. Statistical analyses were performed using linear mixed-effects models fitted to log_10_-transformed values, with strain, treatment, and their interaction as fixed effects and rabbit as a random intercept. Boxes show the median and IQR; whiskers extend to the smallest and largest observations within 1.5 × IQR of the lower and upper quartiles, respectively. \*\*\**P* < 0.001; ns, not significant.

## Discussion

This study identifies bacterial factors important during *S. flexneri* infection *in vivo* and shows that their requirements are shaped by the tissue and temporal contexts. The Tn-seq screen identified 47 canonical pINV-encoded virulence factors while also revealing a broad set of 110 chromosomal fitness factors, whose functions extend beyond the known pINV-centric virulence mechanisms. Experimental analyses of selected candidates *cpxR*, *amiA*, and *zitB* further showed that chromosomal genes can influence distinct stages of infection, from dissemination in epithelial cells to fitness during interactions with host immune cells. The characterization of the role of *zitB* provides a clear example of this contextual fitness dependence: its loss produced no detectable defect during culture media growth, epithelial cell-to-cell spread or early infection *in vivo*, yet the Δ*zitB* mutant showed reduced bacterial burden at 24 hpi in a *MARCO*-expressing macrophage-dependent manner. These findings demonstrate that the importance of a given bacterial gene depends on where and when bacteria encounter particular host pressures.

The extensive identification of pINV gene hits demonstrates both the validity of the screen and the importance of plasmid-dependent virulence mechanisms during intestinal infection. pINV-encoded virulence determinants have been characterized extensively through epithelial culture systems and other experimentally tractable models^55,57,58^. Their depletion in the infant rabbit model confirms that these processes are central during infection of the colonic mucosa *in vivo*. The identification of most genes within the pINV entry region further emphasizes the concerted contribution of this pathogenicity island. The four genes (*ipaJ*, *acp*, *orf131a*, and *orf131b*) at both ends of the entry region were not identified as hits. The functions of Orf131a and Orf131b are poorly understood and may not be required under the condition tested; IpaJ-mediated blockage of intracellular trafficking pathways can be partially compensated by VirA^59^, and the essential chromosomal AcpP (Supplementary Data 1) might provide minimally required compensation for the pINV-encoded Acp. Interestingly, these four genes are conserved in representative plasmid sequences from two major pINV forms, suggesting that their presence may not be completely trivial^60^. In addition, other known pINV virulence factors were also absent from the significant hit set: Serine Protease A (SepA) is known to enhance *S. flexneri* invasion of the intestinal epithelium by disrupting the epithelial barrier^61^. Since SepA acts extracellularly, its activity may benefit neighboring *sepA* mutants during pooled infection in the Tn-seq screen and partially mask the effect of its loss. None of the *osp* or *ipaH* genes was identified as a hit.

Subsets of these effectors target overlapping processes such as NF-κB/MAPK signaling and cell death^34,62^. Disruption of an individual gene may therefore be buffered by the remaining effector repertoire. Nevertheless, the extensive recovery of established pINV virulence genes provides a benchmark for interpreting the less well-characterized chromosomal hits.

The chromosomal hits broaden the framework of *S. flexneri* pathogenesis beyond pINV and demonstrate the importance of preserving both physical and informational integrity: envelope biogenesis and membrane homeostasis maintain the physical structure of the cell, while genome maintenance and translation and protein quality control preserve the transmission and execution of biological information (Fig. 1d). Stress from the host may render defects in either form of integrity consequential for bacterial fitness. Evidently, several hits also have integrity-preserving roles in other pathogens: DsbA-dependent periplasmic protein folding contributes to *Haemophilus influenzae* pathogenesis^63^; HtrA-mediated extracytoplasmic protein quality control maintains the surface and secreted proteome of group B *Streptococcus*; and MnmG–MnmE-dependent tRNA modification supports *Salmonella* Typhimurium virulence^64^. The COG “function unknown” class was the second largest category among the chromosomal hits (Fig. 1d). Upon manual review, the functions of some proteins in this category could be reasonably inferred from the literature, such as the LPS-core kinase WaaY, the cell-shape protein RodZ and the ribosome-splitting factor HflX^40,65,66^. Conversely, the functions of YeeT, the lipoprotein YgdR, and the membrane protein YhdT remain mysterious. A common feature among these proteins is a short, less than 100 predicted amino-acid sequence. Similarly, *yjjY*, one of the strongest hits (log_2_ fold change, −9.63; Holm-adjusted *P* = 9.52 × 10^−8^), received no COG assignment and encodes a predicted protein of 46 amino-acids with unknown functions. Its proximity to the essential *arcA* locus (Supplementary Data 1) raises the question of whether this discovery reflects a requirement for YjjY itself or a local genetic effect on *arcA*.

The phenotypes of *cpxR* and *amiA* illustrate two modes in which chromosomal genes can influence epithelial dissemination. CpxR links chromosomal regulation to the pINV virulence factors through activation of the master regulator *virF* and consequently the downstream cascade in *S. sonnei*^38^. AmiA, by contrast, executes a cell wall peptidoglycan degradation and remodeling function whose importance is selectively revealed in the host cell environment. In organisms that encode a single septal peptidoglycan amidase, such as *Helicobacter pylori*, disruption of *amiA* produces pronounced chaining and defective daughter cell separation^67^. *S. flexneri* and *Escherichia coli* encode three septal peptidoglycan amidases, AmiA, AmiB, and AmiC. In *E. coli*, their activities are partially redundant: loss of a single amidase yields no or mild and low frequency chaining; whereas disruption of all three amidases gives rise to long chaining and pronounced cell separation defect^39^. Consistent with this pattern, phase-contrast microscopy revealed no discernable chaining in either WT or Δ*amiA S. flexneri* during logarithmic or stationary phase growth in LB. Nevertheless, the three amidases differ in their activation and localization. For instance, in *E. coli*, AmiA is diffusely distributed in the periplasm, while AmiB and AmiC are recruited to the division site^68^. In our screen, *amiB* and *amiC* were reduced in the output population but did not meet the output/input ratio threshold of log_2_ fold change −1.585 (*amiB*, locus tag S4592, log_2_ fold change = −0.96; *amiC*, locus tag S3025, log_2_ fold change = −1.54). *amiA* was reduced to a greater extent in the screen (log_2_ fold change = −4.66) and the loss of *amiA* alone resulted in elongated, chain-like structures during bacterial spread in HT-29 cells. Perhaps the epithelial dissemination process imposes demands on cell separation for AmiA that do not inhere in culture media growth. Because cell-to-cell spread involves actin-based motility, membrane protrusion formation and resolution, and escape from double-membrane vacuoles^13^, inefficient separation could interfere with these steps, although the exact mechanism remains to be explored.

The *zitB* phenotype provides insight into how innate immunity curbs *S. flexneri*. Macrophage depletion eliminated the burden difference between WT and Δ*zitB*, even when the *LCN2*-expressing neutrophil response remained robust. Together with the impaired survival of Δ*zitB* in THP-1-derived macrophages, these findings support a model in which ZitB helps *S. flexneri* withstand a macrophage-induced pressure (Fig. 7). This conclusion complements a recent study in a *Nlrc4*^−/–^*Casp11*^−/–^ murine model showing that while neutrophils were dispensable for controlling *Shigella*, macrophages resolved infection indirectly through TLR-induced IL-12 production and downstream IFN-γ-mediated control of bacteria in intestinal epithelial cells^69^. Our findings suggest that macrophages may also impose pressures directly, resulting in specific requirement for bacterial fitness factors in non-epithelial niches.

**Fig. 7:**
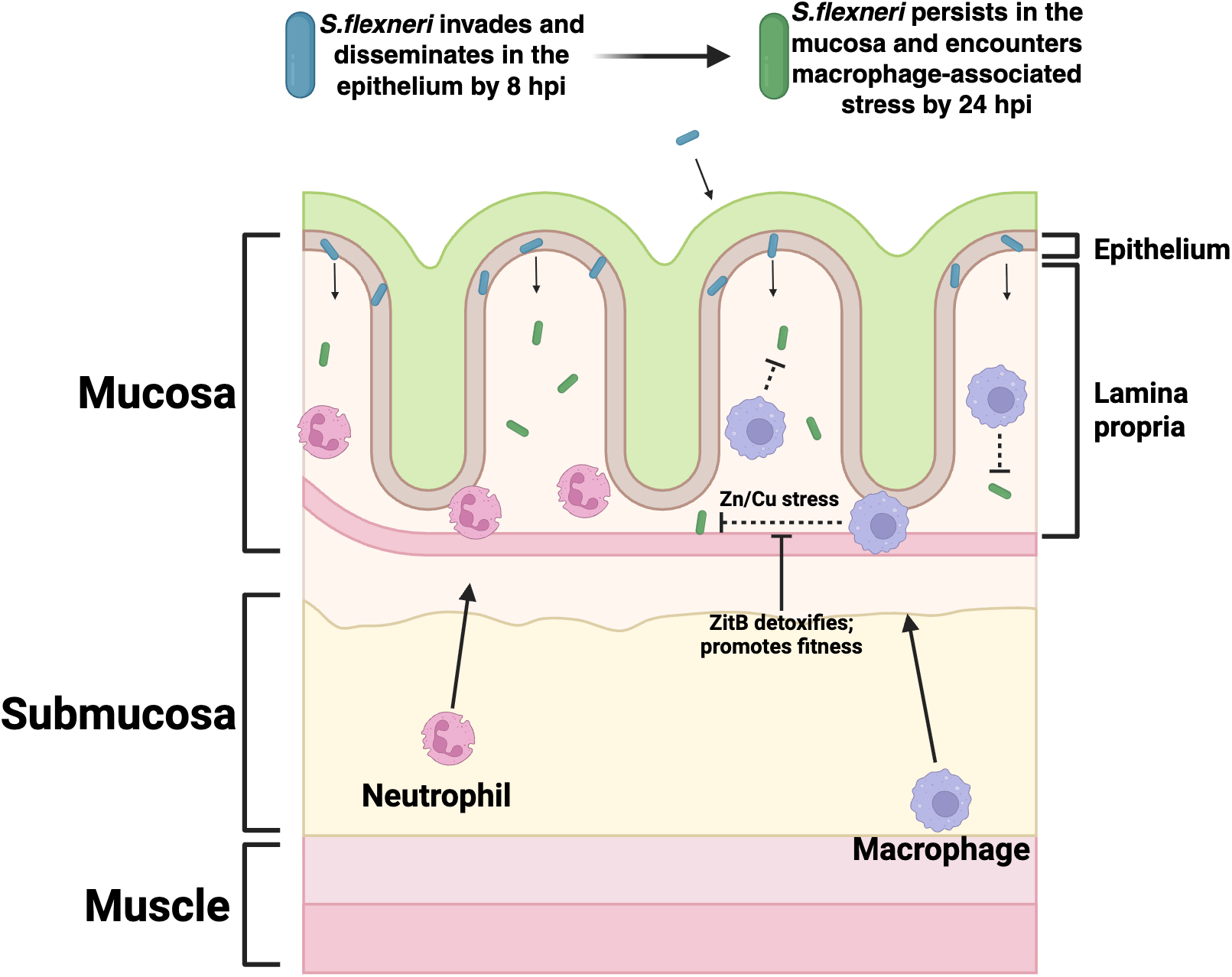
Proposed model for contribution of ZitB to *S. flexneri* fitness *in vivo*. By 8 hpi, *S. flexneri* invades and disseminates within the colonic epithelium. By 24 hpi, bacteria persist within the mucosa and encounter macrophage-associated stress. The data support a model in which ZitB promotes *S. flexneri* fitness by detoxifying Zn/Cu-associated stress imposed by macrophages. Neutrophil responses remain prominent but does not explain the ZitB-dependent bacterial burden difference. Blue and green bacteria denote the earlier and later stages of infection, respectively. This model is schematic and does not represent the relative sizes or numbers of bacteria, neutrophils, or macrophages.

Metal intoxication provides a plausible link between ZitB and resistance to macrophage-induced stress. Loss of *zitB* increased susceptibility to zinc and copper in minimal medium, and complementation restored full growth, while TPEN eliminated the detectable difference between WT and Δ*zitB* during infection of THP-1-derived macrophages. Both zinc and copper have been implicated as antimicrobial agents against bacterial pathogens deployed by macrophages^53,54^.

Thus, the sensitivity of Δ*zitB* to both metals and the rescue of its macrophage-associated defect following TPEN treatment point to zinc, copper, or a combination of both. Why ZitB was required despite the presence of the additional main zinc and copper exporters ZntA and CopA remains unclear. Loss of *zitB* alone sufficiently increased sensitivity to both zinc and copper in M9 media; *zitB* alone, and not *zntA* or *copA*, was identified as a hit in the Tn-seq screen (*zitB*, locus tag S0560, log_2_ fold change = −5.17; *zntA*, locus tag S4276, log_2_ fold change = 0.68; *copA*, locus tag S0436, log_2_ fold change = 0.23) (Supplementary Data 1 and Supplementary Data 2). ZitB, being a CDF family transporter, differs from these ATP-dependent exporters in both transport mechanism and regulation^70–72^, which may allow it to operate over different dynamic range and temporal pattern in response to excessive metal ions. Additionally, disruption of ZitB may perturb both zinc and copper homeostasis and cause synergistic intoxication.

Despite advancing the understanding of *S. flexneri* pathogenesis, this study has several limitations. Tn-seq is limited by population bottlenecks that can cause stochastic loss. Intrarectal inoculation alleviated this problem by focusing the screen on direct colonic infection. But this design was accompanied by a tradeoff for a limitation in scope, wherein genes required for survival and dissemination through the upper gastrointestinal tract, transient colonization or expansion in intermediate niches, biofilm formation^73^, or interactions with the microbiota would not be identified. This study can also be furthered by expanding the temporal and immunological perspectives of infection. The 8 and 24-hpi time points do not capture the later resolution phase, and *MARCO* and *LCN2* RNAscope analysis provides an incomplete picture that does not fully depict immune cell populations, their activation states, or cytokine responses. Future work in transcriptome and metabolome analyses could provide systemic perspectives of the host environment.

*Shigella* is a highly human-adapted pathogen. Through its convergent evolutionary history, it has diverged from commensal *E. coli* in both gain and loss of gene functions, such as the acquisitions of the pINV and O antigen gene clusters and the losses of flagellar system and metabolic flexibility^74,75^. Consistent with this highly developed adaptation, our screen did not identify insertions whose disruption significantly increased fitness *in vivo* (Supplementary Data 2). Because humans are the only known natural reservoir for *Shigella*, transmission, colonization, infection and sustaining infection in humans must be vital for *Shigella* to maintain its biomass in the biosphere. As a result, such a lifestyle would require adaptation to the distinct physiological niches encountered within the human host. *zitB* represents a chromosomal fitness factor whose contribution emerges only during interaction with immune cells, particularly macrophages. ZitB most plausibly promotes fitness by helping *S. flexneri* withstand metal intoxication stresses imposed within or by the macrophage niche (Fig. 7). More broadly, this study signified the utility of a framework for mapping bacterial fitness determinants to the temporal stages of infection, and spatial and cellular niches. Such a “fitness-factor atlas” can refine pathogenies understanding and guide rational drug design.

## Methods

### Bacterial strains and culture conditions

The wild-type (WT) strain used in this study was *Shigella flexneri* serotype 2a strain 2457T^10^. The Δ*zitB*, Δ*cpxR*, Δ*amiA*, Δ*waaY*, Δ*hslV*, and Δ*hha* allelic-exchange mutants were generated by replacing each corresponding coding sequence with a kanamycin-resistance cassette through homologous recombination using the suicide vector pSB890, as previously described^73^. The Δ*zitB* complementation strain (Δ*zitB*-comp) was generated by expressing *zitB* under its native promoter in the pBAD18 vector (ATCC 87393). CFP-expressing WT and Δ*zitB* strains were generated by introducing pMMB207, a chloramphenicol-resistance plasmid carrying the gene encoding cyan fluorescent protein (CFP) under the control of an isopropyl β-D-1-thiogalactopyranoside (IPTG)-inducible promoter. pBAD18 and pMMB207 were introduced by electroporation. Unless otherwise indicated, bacteria were first cultured on LB agar containing 100 μg/ml Congo red. Bacterial strains were then cultured in lysogeny broth (LB) at 37 °C with aeration. For preparation of exponential phase cultures for HT-29 and THP-1 macrophage infection experiments, the preceding overnight cultures were grown in LB at 30 °C with aeration. For infant rabbit infections, bacterial strains were cultured in tryptic soy broth (TSB) at 37 °C for 12.5 h with aeration. For growth curve experiments in rich media, overnight cultures grown at 30 °C were washed in Dulbecco’s phosphate-buffered saline (DPBS) and diluted into the indicated medium to a starting optical density at 600 nm (OD_600_) of 0.01. For growth measurements in LB, 5 ml cultures were incubated in standard culture tubes at 37 °C with aeration, and OD_600_ was measured manually at the indicated time points using a GENESYS 20 spectrophotometer (Thermo Scientific). For growth measurements in TSB, bacteria were cultured in 96-well microplates at 37 °C with orbital shaking, and OD_600_ was measured every 10 min using a SpectraMax iD3s (Molecular Devices). The volume in each well was 200 µl. For growth experiments in M9 minimal medium, M9 medium was prepared by diluting a 5× M9 salts solution (Sigma-Aldrich) with water and supplementing with glucose, nicotinic acid, methionine, and tryptophan to final concentrations of 0.4% (w/v), 12.5 μg/ml, 45 μg/ml, 20 μg/ml, respectively. Individual colony from LB agar was suspended in DPBS and inoculated into M9 medium alone or M9 medium supplemented with the indicated metal salts. Cultures were incubated at 37 °C for 16 h with aeration. Metal salts and supplier were: zinc sulfate heptahydrate (BioReagent grade), copper(II) sulfate pentahydrate (99.999% trace metals basis), iron(II) sulfate heptahydrate (BioReagent grade), manganese(II) sulfate monohydrate (ReagentPlus grade), all from Sigma-Aldrich. *E. coli* strains SM10λpir, Δ*nic*35 and DH5α were used for cloning purposes. 2,6-diaminopimelic acid (DAP) was supplemented (100 μg/ml) for the growth of *E. coli* strain Δ*nic*35. Primers used in this study are listed in Supplementary Table 1. Antibiotics were added as required for strain selection or plasmid maintenance at the following concentrations: ampicillin (100 µg/ml), chloramphenicol (10 µg/ml), kanamycin (30 µg/ml) or tetracycline (10 µg/ml).

### Transposon sequencing screen and data analysis

A saturated mini-Tn10 (mTn10) insertion library was generated essentially as described^76^. Briefly, transposition plasmid pDL1093 was moved into *S. flexneri* 2457T by mating and propagated in LB or LB agar supplemented with 5 µg/ml chloramphenicol and 50 µg/ml kanamycin sulfate at the permissive temperature for plasmid replication, 30 °C. In duplicate, 20 fresh colonies were pooled and used to inoculate 30 ml of LB broth supplemented with the same antibiotics in 50-ml Erlenmeyer flasks and incubated overnight at 30 °C with aeration. The next morning, 20 µl of each overnight culture was added to 100 ml of LB broth supplemented with 100 µg/ml kanamycin sulfate in 250-ml Erlenmeyer flasks and incubated for 8 h at 40 °C with aeration to induce transposition and halt plasmid replication. The resulting early stationary-phase cultures were used to seed a second passage by adding 0.25 ml of each culture to new flasks containing 100 ml LB supplemented with kanamycin sulfate and incubating overnight at 40 °C with aeration. This second passage ensured the elimination of cells lacking mTn10 insertions. The two final cultures were pooled and cryopreserved in multiple single-use 1-ml aliquots after addition of glycerol to 20% (v/v). For *in vivo* Tn-seq, four aliquots of the transposon library were thawed and grown in TSB for 12.5 h. The overnight cultures were collected and resuspended in 200 μl PBS per animal and used to infect four infant rabbits. At 24 hpi, colons were collected, homogenized in 10 ml PBS, and homogenates were inoculated into 500 ml LB with 200 µg/ml kanamycin for outgrowth before DNA extraction. Transposon junction sequencing was performed essentially as described^76^. Briefly, total DNA was isolated from the input and each output using the DNeasy Blood & Tissue Kit (Qiagen) according to the manufacturer’s instructions for Gram-negative bacteria, including Proteinase K treatment. Next, the mTn10 junctions were amplified from the input library and each output library by HTML-PCR followed by sequencing on the Illumina platform. Reads were trimmed based on quality and poly-C or poly-G sequence at the 3′ ends were removed. The trimmed reads were mapped separately to the *S. flexneri* chromosome and pINV, and DvalGenome values were calculated for each gene using the custom Python script Hopcount^76^. Only uniquely mapping reads were analyzed; therefore, repeated sequences, such as ribosomal operons or natural transposons, lack mapped reads. Essential genes were defined as non-repeated genes with DvalGenome values <0.01 in the input library and were manually confirmed by visually examining the pattern of insertions in and surrounding each gene using a genome browser as described^76^.

### COG annotation and STRING network analysis

Protein sequences corresponding to the 110 significant chromosomal hits were retrieved from KEGG^77^ using the KEGGREST package in R and analyzed with eggNOG-mapper (version 5.0.2)^78^ on usegalaxy.eu. COG assignments were extracted from the annotation output and matched to COG2024 functional categories^79^. Genes assigned to multiple categories were counted in each category. The chromosomal hits were also analyzed using STRING^80^. Interactions with a minimum interaction score of 0.7 were retained, and network clusters were identified using the Markov Cluster Algorithm with an inflation parameter of 3. Gene level annotations, category definitions, STRING mappings, interactions and cluster assignments are provided in Supplementary Data 3.

### HT-29 cell culture and dissemination assays

HT-29 (ATCC) epithelial dissemination assays were performed as previously described^37^. Confluent HT-29 monolayers in 96-well plates in 100 µl McCoy’s 5A medium with 10% heat-inactivated fetal bovine serum (HI-FBS) per well were infected with exponential phase *S. flexneri*. Equal volume dose was used for all strains by diluting 75 µl exponential phase bacterial culture into 6 ml McCoy’s 5A medium with FBS, and from which 50 µl was subsequently added to each well. Plates were centrifuged at 130 × g for 5 min at room temperature and incubated for 1 h at 37 °C with 5% CO_2_. The plates were then supplemented with 50 µl McCoy’s 5A medium with 10% HI-FBS containing gentamicin (50 µg/ml) and IPTG (10 mM) for each well to eliminate extracellular bacteria and induce CFP expression, respectively. At 8 hpi, cells were fixed with 4% paraformaldehyde, and infection foci were detected and quantified using an ImageXpress Micro imaging system and MetaXpress software (Molecular Devices).

### Animal procedures

Infant rabbit infections were performed as previously described^23^. Pregnant New Zealand White rabbits were obtained from Charles River Laboratories. Newborn rabbits were either housed with the dam until experimental procedures or isolated after birth and maintained in a 30 °C incubator. For infection, 10–15-day-old infant rabbits were infected intrarectally using 20-gauge sterile feeding tubes (Instech Labs) under anesthesia with 5% isoflurane in 5 l/min oxygen. 5 ml culture was collected and resuspended in 200 μl PBS for each infant rabbit (∼10^9^ CFU per animal). At the indicated time points, animals were euthanized according to approved procedures, and colons were collected for histological analysis, or homogenized in sterile PBS, diluted and subsequently plated on selective agar media to determine CFU and competitive index. Clinical signs of blood and diarrhea were scored blindly at the time of tissue harvest based on the color, wetness and anatomical extent of staining on the hindlimb fur using the following scale: 0, no detectable signs; 1, staining restricted to the genital region; 2, staining extending to the genital region and abdomen; and 3, staining extending to the genital region, abdomen and legs. For macrophage depletion experiments, Clodrosome (500 µl per animal) containing active clodronate was administered through the lateral saphenous vein 48 h before infection, with Encapsome used as the corresponding control (Encapsula NanoSciences). All BSL2 and animal procedures were reviewed and approved by the University of Virginia Institutional Biosafety Committee and the Institutional Animal Care and Use Committee under protocols 3923-15 and 4161, respectively.

### Tissue processing, staining and quantitative image analysis

Following euthanasia, a 10-cm segment of the distal colon was harvested and processed for paraffin embedding. Colons were rinsed with PBS, flushed with modified Bouin’s fixative (50% ethanol, 5% glacial acetic acid, and 45% distilled water), opened longitudinally, and arranged as Swiss rolls in tissue cassettes. Cassettes were immersed in neutral-buffered formalin for 48 h, after which tissues were transferred to 70% ethanol and processed for dehydration and paraffin infiltration. Tissues were manually embedded in paraffin, and 5-μm sections were cut using a Leica microtome. Paraffin sections were stained with hematoxylin and eosin (H&E) and imaged using an Aperio ScanScope whole-slide scanner (Leica Biosystems). Epithelial fenestration was measured along the evaluable colon section using Aperio ImageScope. The percentage of epithelial fenestration was calculated for each colon as the length of mucosa exhibiting epithelial fenestration divided by the total evaluated mucosal length, multiplied by 100.

Immunofluorescence staining of paraffin-embedded sections was performed as previously described^36^ with modifications. Sections were deparaffinized in xylene and rehydrated through graded ethanol solutions into water. Heat-induced antigen retrieval was performed in citrate-based antigen-retrieval buffer (Vector Laboratories, H-3300) using an electric pressure cooker (Instant Pot) at high pressure for 1 min, followed by rapid depressurization. Sections were permeabilized with 0.1% Triton X-100, blocked in PBS containing 5% bovine serum albumin and 2% normal goat serum, and incubated overnight at 4 °C with antibodies against E-cadherin (1:100; BD Biosciences, 610181) and *Shigella* spp. (1:100; ViroStat, 0901). After washing with PBS, sections were incubated with goat anti-mouse IgG and goat anti-rabbit IgG secondary antibodies (1:500; Thermo Fisher Scientific) conjugated to Alexa Fluor 568 (A-11004) and 488 (A-11034), respectively, or Alexa Fluor 594 (A-11032) and 514 (A-31558), respectively, depending on channel assignment. After incubation with secondary antibodies for 2 h at room temperature, sections were mounted using ProLong Gold Antifade Mountant (Thermo Fisher Scientific). RNAscope was performed on paraffin-embedded sections using the RNAscope Multiplex Fluorescent Reagent Kit v2 (Advanced Cell Diagnostics) according to the manufacturer’s instructions. Briefly, following deparaffinization, hydrogen peroxide treatment, target retrieval, and protease treatment, sections were hybridized with probes targeting rabbit *MARCO* (Advanced Cell Diagnostics, 847591) or *LCN2* (Advanced Cell Diagnostics, 857241).

Each probe was assigned to channel C1; channels C2 and C3 were not used. Signals were amplified and developed using the C1 channel with Opal 690 fluorophore (1:1000; Akoya Biosciences, FP1497001KT). If applicable, sections were subsequently immunostained as described above. All sections were then counterstained with DAPI and mounted using ProLong Gold Antifade Mountant. Immunofluorescence and RNAscope stained sections were imaged using either an ImageXpress Micro imaging system or a Nikon TE2000 microscope equipped with a Hamamatsu ORCA-ER CCD camera, and images were analyzed using MetaXpress software (Molecular Devices). The image acquisition and quantification workflow is summarized in Supplementary Fig. 5. Signal positive area was identified by image thresholding, and the relevant anatomical region was delineated within each acquired image. *S. flexneri* positive area was normalized to mucosal area, whereas *MARCO* and *LCN2* positive areas were normalized to the corresponding analyzed area of the mucosa, submucosa or muscle. Normalized measurements were reported per 10,000 px^2^ of anatomical area. To assess whether positive area provided a representative measure of marker abundance, normalized *MARCO* positive area was compared with normalized *MARCO* object count in a validation subset of images (Supplementary Fig. 5f). Multiple regions were quantified from each rabbit and analyzed as nested observations using linear mixed-effects models.

### THP-1-derived macrophage infection and cytotoxicity assays

THP-1 cells (ATCC) were cultured in RPMI 1640 containing 10% HI-FBS. Differentiation was induced by 200 nM phorbol-12-myristate-13-acetate (PMA, Sigma-Aldrich) for 48 hours, followed by an additional 48-hour rest period in fresh medium in 24-well plates with coverslips added. For infection, each well was seeded with 500 μl medium containing 4 × 10^5^ cells/ml THP-1 cells. For inoculation, fresh media containing bacteria were added. Exponential phase cultures of *S. flexneri* were washed by DPBS and diluted to RPMI 1640 medium with 10% HI-FBS for a multiplicity of infection (MOI) of 2. When applicable, N,N,N′,N′-Tetrakis(2-pyridylmethyl)ethylenediamine (TPEN, Sigma-Aldrich) was added to a final concentration of 10 μM. Plates were centrifuged at 130 × g for 10 min at room temperature and incubated for 1 h at 37 °C with 5% CO_2_, after which gentamicin (50 µg/ml) was added to eliminate extracellular bacteria. IPTG (10 mM) was added to induce CFP expression for infection with CFP-expressing bacteria and imaging-based analysis. At indicated time points, coverslips were fixed in 4% paraformaldehyde in a separate plate, and immunostaining was performed for *S. flexneri* as follows: coverslips were permeabilized with 0.1% Triton X-100, blocked in PBS containing 5% bovine serum albumin and 2% normal goat serum, and incubated overnight at 4 °C with anti-*Shigella* spp. antibody (1:100; ViroStat, 0901). After PBS washing, coverslips were incubated with goat anti-rabbit IgG Alexa Fluor 514 or 647 (1:500; Thermo Fisher Scientific, A-31558/A-21244) secondary antibody for 2 h at room temperature and mounted using DABCO-based antifade mounting medium. For CFU enumeration, coverslips were permeabilized with 0.1% Triton X-100 for 10 min, mechanically scraped with a P1000 pipette tip, and the lysates were plated on selective agar media. For quantification of CFP-expressing *S. flexneri* associated with macrophages, images were acquired using an ImageXpress Micro imaging system or a Nikon TE2000 microscope equipped with a Hamamatsu ORCA-ER CCD camera and quantified using the Multi-Wavelength Cell Scoring function in MetaXpress software (Molecular Devices). CFP-positive signal was expressed as a percentage of the total *Shigella* signal identified by immunostaining. Lactate dehydrogenase (LDH) release was measured using the CytoTox 96 Non-Radioactive Cytotoxicity Assay (Promega, G1780) according to the manufacturer’s instructions. Culture supernatants (30 μl) were transferred to a clear 96-well plate, incubated with an equal volume of CytoTox 96 reagent for 30 min at room temperature protected from light, and the reaction was stopped using the supplied Stop Solution. Absorbance was measured at 490 nm using a microplate reader (PerkinElmer VICTOR^3^ 1420). Percent cytotoxicity was calculated after correction for spontaneous release from uninfected cells (Supplementary Fig. 8i) and normalization to maximum LDH release by cell lysis.

### Statistical analysis and data visualization

Statistical analyses and data visualization were performed in R (4.6.1)^81^ using RStudio (2026.7.0.139). Data processing was performed using the tidyverse package^82^. Unless otherwise specified, continuous outcomes were analyzed using linear models. Estimated marginal means and post hoc contrasts were obtained using the emmeans package^83^, with Tukey adjustment for pairwise comparisons and Dunnett adjustment for comparisons to WT. Model predicted values displayed in the figures were generated using the ggeffects package^84^. Colonic bacterial burden and RNAscope marker measurements obtained from multiple quantified regions within each rabbit were analyzed using linear mixed-effects models fitted with lme4 and lmerTest^85,86^. Unless otherwise specified, strain was included as a fixed effect and rabbit as a random intercept. For experiments involving both strain and treatment, additive models were compared with models containing the strain × treatment interaction term.

Likelihood-ratio test, Akaike information criterion (AIC) and Bayesian information criterion (BIC) were used to evaluate model fit. The interaction term was retained when supported by the model comparisons. Model assumptions were evaluated by inspection of residual-versus-fitted and normal quantile–quantile plots. Outcomes were log_10_-transformed when necessary to improve normality. Unless otherwise stated, statistical tests were two-sided, and *P* < 0.05 was considered statistically significant. Data were visualized using ggplot2^87^, ggprism, and ggbeeswarm. Fig. 7 was created with BioRender.com.

## Data availability

The sequencing data generated in this study will be deposited in the NCBI Sequence Read Archive (SRA), and the accession number will be provided upon availability. All remaining data from this study are provided in the Supplementary Information and Source Data file.

## Acknowledgments

We thank all the members of the Agaisse laboratory for the discussions on the manuscript. We thank Jeremy Gatesman and Deanna Zielinski for veterinary and technical assistance with clodronate liposome administration. We thank Healy Rosenberg, Hannah Collier, Todd Kirks, Alice Kweon, Zackary Lifschin, Liam Myers and Aaron Tang for assistance with the care and nursing of infant rabbits. We also thank Sheri Vanhoose and the UVA Research Histology Core for preparing paraffin sections. This work was supported by the National Institutes of Health grants R01AI073904, R01AI179778 (H.A.) and R21AI178051 (H.A. and A.C.).

## Author information

### Contributions

Y.S. and H.A. conceived and designed the study. A.C. constructed the Tn-seq library, L.K.Y. conducted the Tn-seq screen, A.C. conducted the bioinformatic analysis of the sequencing results and H.A. constructed the validation mutant strains. Y.S. conducted all *in vitro* and *in vivo* experiments. Y.S., H.A., D.W.L., and A.C. analyzed the data. Y.S. and H.A. wrote the paper.

Corresponding author Correspondence to Hervé Agaisse.

## Ethics declarations

### Competing interests

The authors declare no competing interests.

## Supplementary Information

**Supplementary Fig 1.**
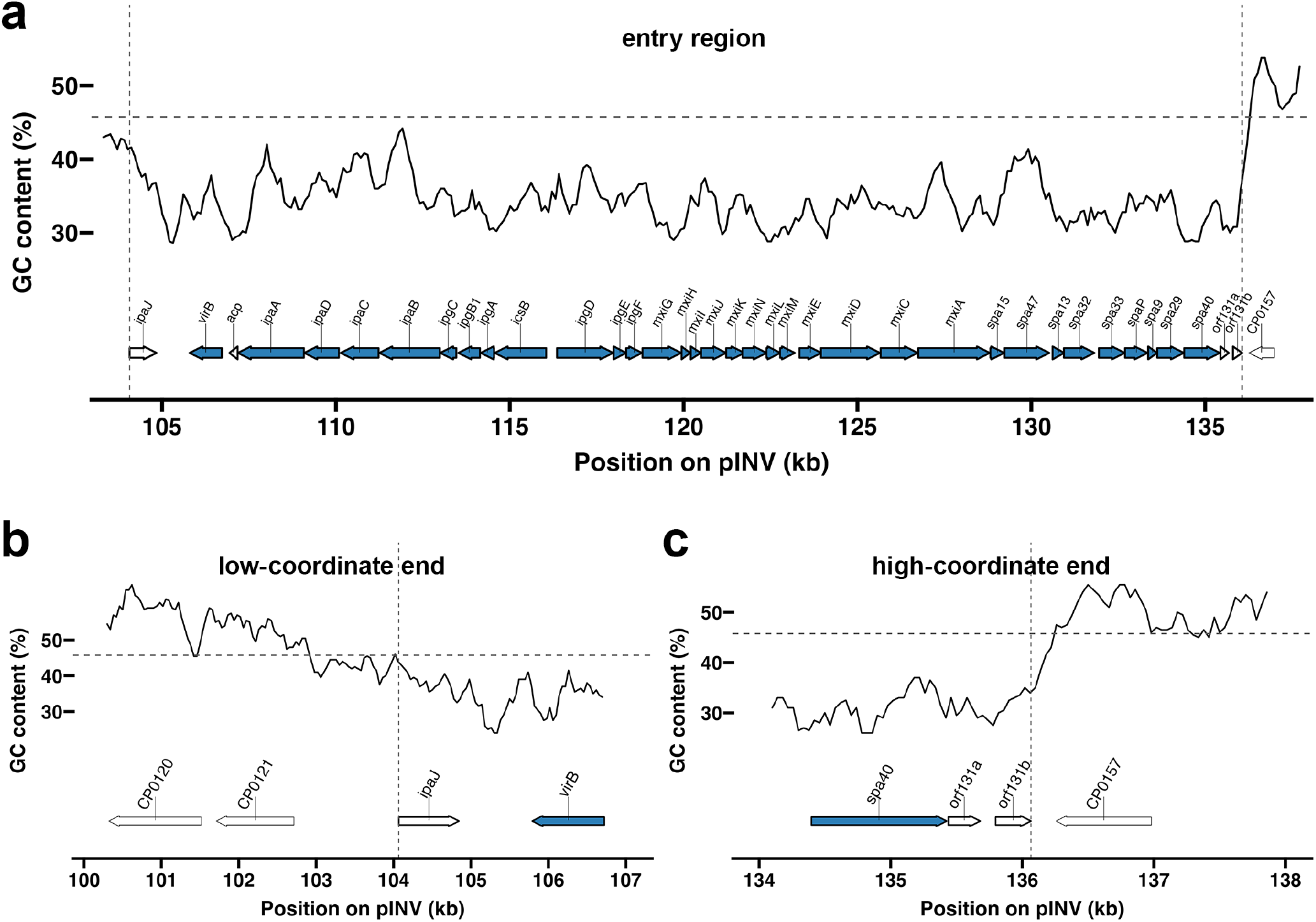
GC content across the pINV entry region and its flanking sequences. GC content is plotted against pINV coordinates using sliding-window analysis, with gene organization shown beneath each plot. Gene arrows corresponding to loci within the pINV entry region have bold outlines, and genes identified as hits in the Tn-seq screen are shown in blue. Vertical dashed lines mark the boundaries of the pINV entry region, and the horizontal dashed line indicates the mean GC content of the entire pINV plasmid. **a** GC content across the pINV entry region and adjacent flanking sequences, calculated using a 500-bp window and a 100-bp step size. **b** GC content across the low-coordinate end of the pINV entry region and its adjacent flanking sequence, calculated using a 200-bp window and a 40-bp step size. **c** GC content across the high-coordinate end of the pINV entry region and its adjacent flanking sequence, calculated using a 200-bp window and a 40-bp step size.

**Supplementary Fig. 2:**
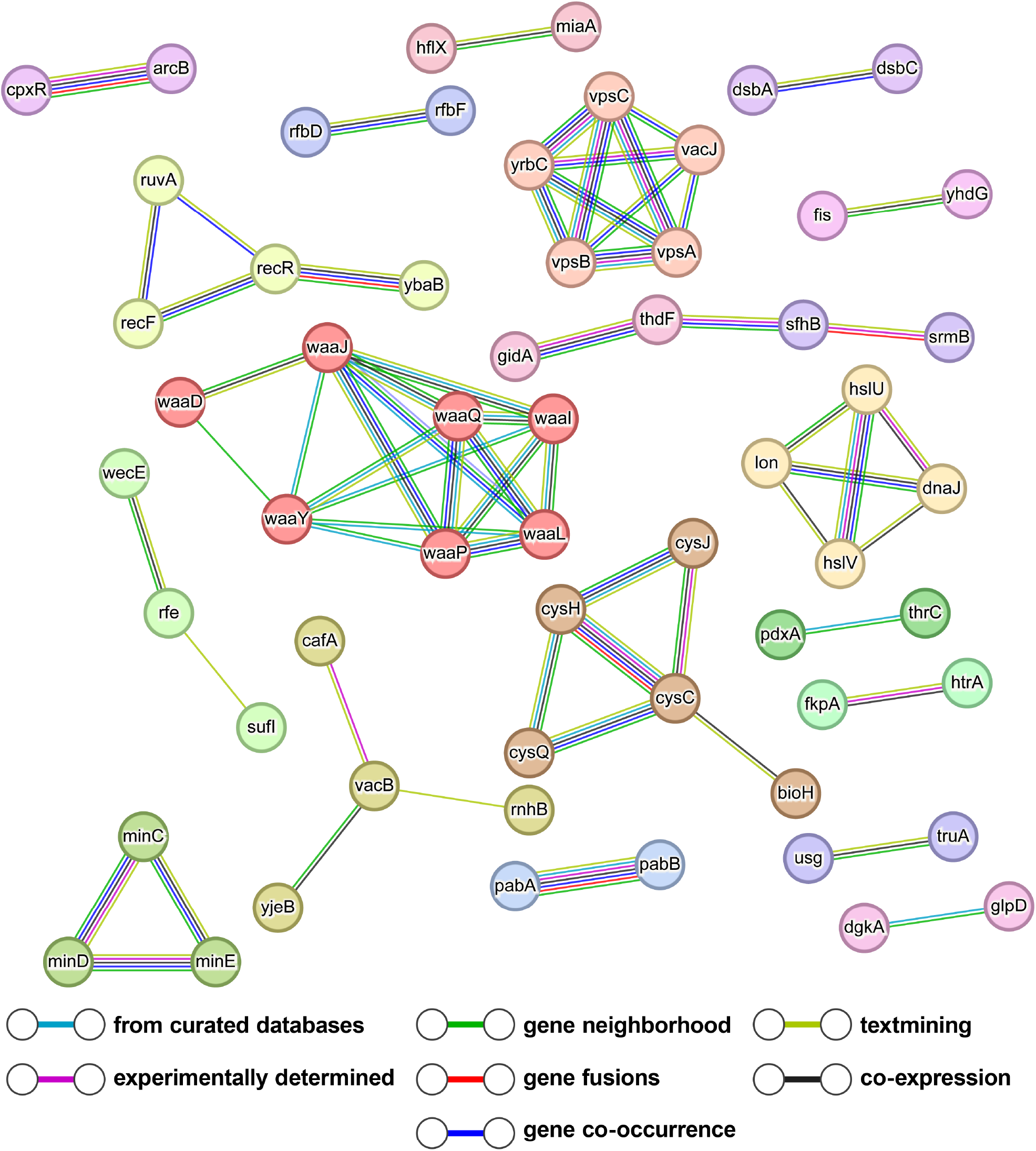
Functional association network of chromosomal Tn-seq hits. Chromosome-encoded Tn-seq hits were analyzed using STRING with a minimum required interaction score of 0.7. Nodes represent proteins and are colored according to Markov cluster algorithm clusters generated using an inflation parameter of 3. Edges represent known or predicted functional associations, with line colors indicating the supporting evidence sources shown in the key. Disconnected nodes are not shown.

**Supplementary Fig. 3:**
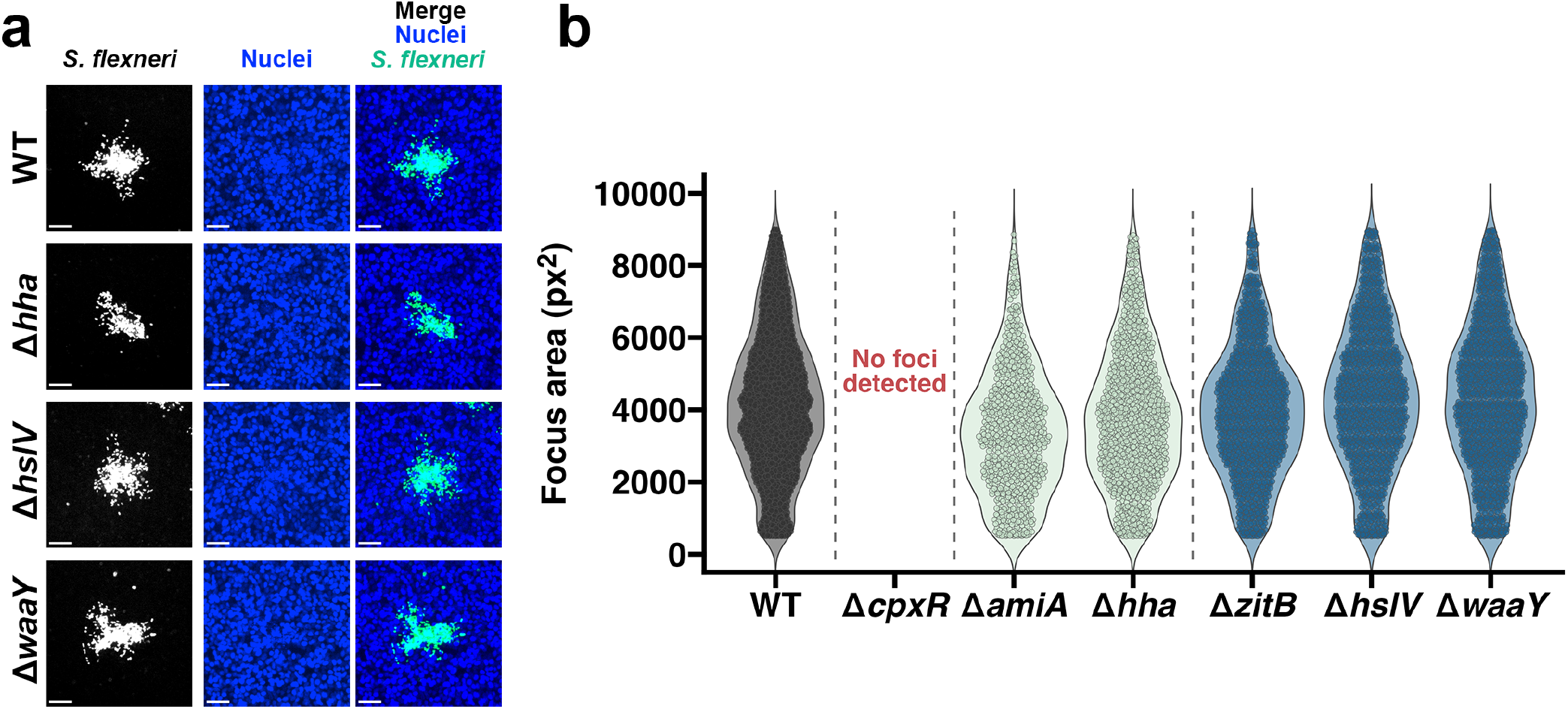
Additional cell-to-cell spread analysis of selected Tn-seq candidate mutants in HT-29 cells. **a** Representative images of infection foci formed by WT, Δ*hha*, Δ*hslV*, and Δ*waaY* in HT-29 cell monolayers at 8 hpi. *S. flexneri* is shown in white in the single-channel images and in green in the merged images; nuclei are shown in blue. Scale bars, 50 μm. **b** Violin plots showing the distributions of individual focus areas formed by WT and the indicated mutants in HT-29 cell monolayers at 8 hpi. Each point represents an individual infection focus pooled across three independent experiments (*n* = 3). No foci were detected for Δ*cpxR*.

**Supplementary Fig. 4:**
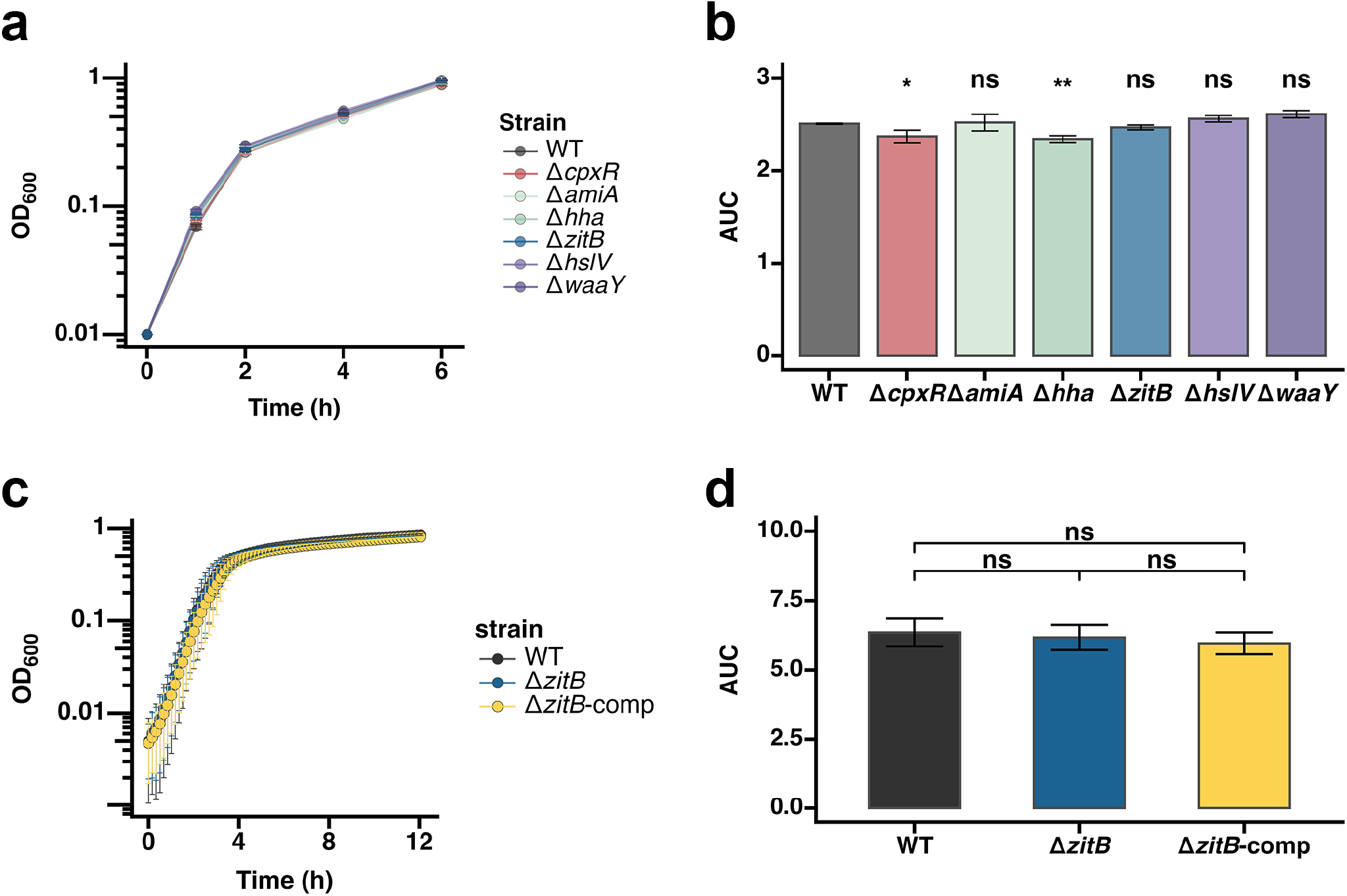
*In vitro* growth of selected strains. **a** Growth of WT and the indicated mutants in LB broth, measured by OD_600_ over 6 h. Data are shown as mean ± SD from three independent cultures (*n* = 3). **b** Area under the curve (AUC) calculated from the OD_600_ measurements in **a**. Each point represents one independent culture, and bars show mean ± SD (*n* = 3). Statistical analysis was performed using a linear model followed by Dunnett-adjusted comparisons of each mutant with WT. **c** Growth of WT, Δ*zitB*, and Δ*zitB*-comp in TSB, measured by OD_600_ over 12 h. Data are shown as mean ± SD from three independent experiments (*n* = 3). **d** AUC calculated from the OD_600_ measurements in **c**. Each point represents one independent experiment, and bars show mean ± SD (*n* = 3). Statistical analysis was performed using a linear model followed by Tukey-adjusted pairwise comparisons. \**P* < 0.05; \*\**P* < 0.01; ns, not significant.

**Supplementary Fig. 5:**
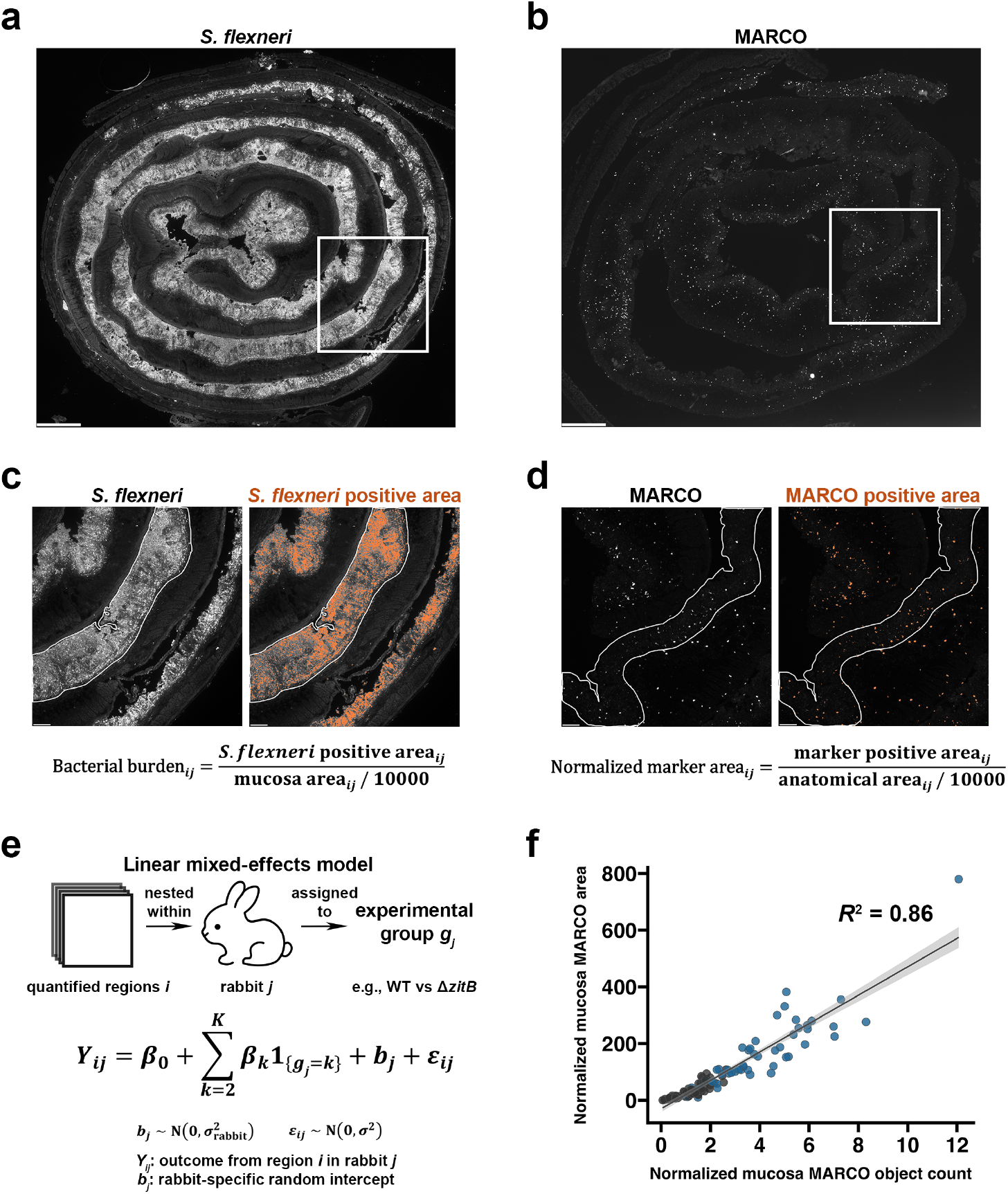
Workflow for tissue image quantification and mixed-effects modeling. **a**–**b** Representative montages of colon Swiss rolls showing *S. flexneri* signal (**a**) and *MARCO* signal as an example of RNAscope marker imaging (**b**). Each montage was assembled from acquired images to provide an overview of the tissue section. White rectangles indicate the acquired images shown in **c** and **d**, respectively. Scale bars, 500 μm. **c**–**d** Acquired images indicated in **a** and **b**, respectively, showing the original *S. flexneri* (**c**) or *MARCO* (**d**) signal and the corresponding thresholded positive area in orange. White lines delineate the mucosa. Signal-positive area and the corresponding anatomical area were measured within each quantified region. Bacterial burden was normalized to mucosal area (**c**), whereas marker-positive area was normalized to the anatomical area analyzed (**d**). Normalized values were reported using a 100 px × 100 px reference area (10,000 px^2^). The marker-quantification workflow was applied to *MARCO* and *LCN2* in the mucosa, submucosa, and muscle. Scale bars, 100 μm. **e** Schematic of the hierarchical analysis. Quantified regions were nested within rabbits, which were assigned to experimental groups, and outcomes were analyzed using linear mixed-effects models with experimental group as a fixed effect and rabbit as a random intercept. **f** Relationship between normalized mucosal *MARCO*-positive area and normalized mucosal *MARCO* object count in a validation subset of 113 quantified regions from rabbits infected with WT or Δ*zitB* rabbits (*n* = 4 rabbits per group). Each point represents one quantified region; gray and blue indicate WT and Δ*zitB*, respectively. The line shows the linear regression fitted to the combined data from both groups; shading indicates the 95% confidence interval (*R*^2^ = 0.86).

**Supplementary Fig. 6:**
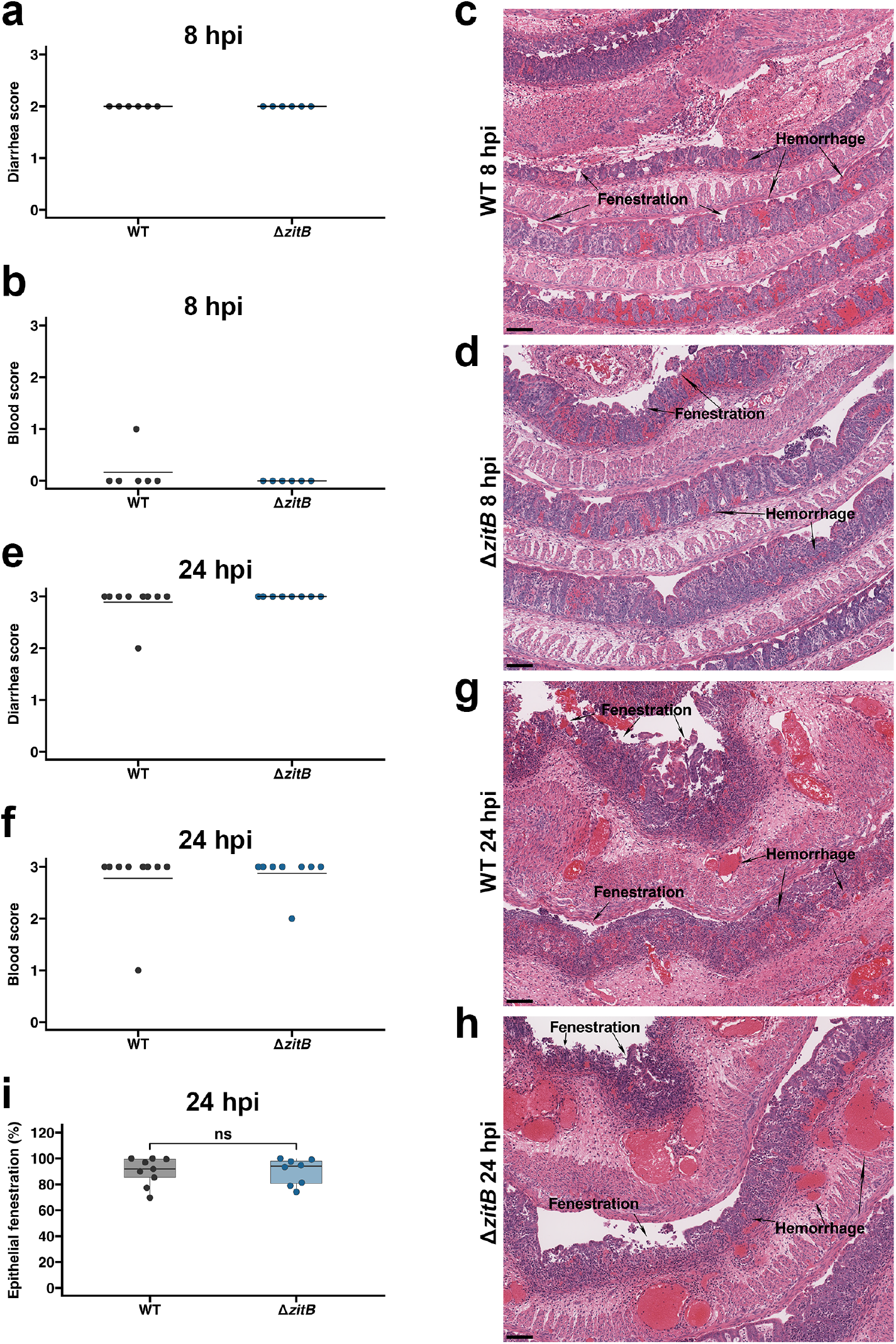
WT and Δ*zitB* infections show comparable intestinal pathology and epithelial fenestration. **a**–**b** Diarrhea (**a**) and blood (**b**) scores in rabbits infected with WT or Δ*zitB* at 8 hpi. Each point represents one rabbit, and horizontal lines indicate the mean (*n* = 6 per group). **c**–**d** Representative hematoxylin and eosin (H&E)-stained colon sections from rabbits infected with WT (**c**) or Δ*zitB* (**d**) at 8 hpi. Arrows indicate examples of epithelial fenestration and hemorrhage. Scale bars, 100 μm. **e**–**f** Diarrhea (**e**) and blood (**f**) scores in rabbits infected with WT or Δ*zitB* at 24 hpi. Each point represents one rabbit, and horizontal lines indicate the mean (*n* = 9 for WT and *n* = 8 for Δ*zitB*). **g**–**h** Representative H&E-stained colon sections from rabbits infected with WT (**g**) or Δ*zitB* (**h**) at 24 hpi. Arrows indicate examples of epithelial fenestration and hemorrhage. Scale bars, 100 μm. **i** Percentage of the traced colonic epithelial length exhibiting fenestration at 24 hpi. Each point represents one rabbit (*n* = 9 for WT and *n* = 8 for Δ*zitB*). Boxes show the median and interquartile range (IQR); whiskers extend to the smallest and largest observations within 1.5 × IQR of the lower and upper quartiles, respectively. Statistical analysis was performed using a linear model. ns, not significant.

**Supplementary Fig. 7:**
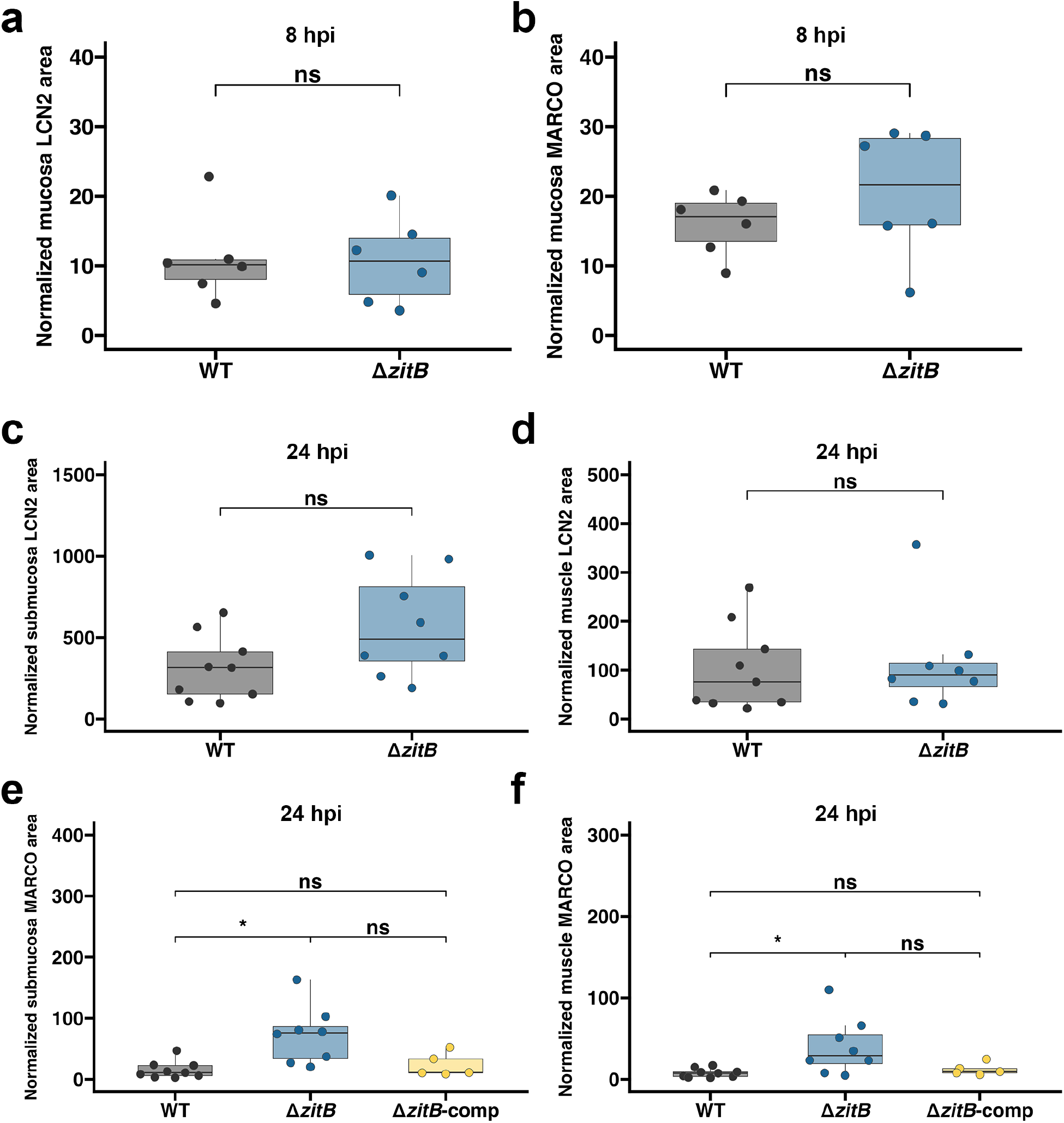
Additional analyses of *LCN2* and *MARCO* across intestinal tissue compartments. **a**–**b** Normalized mucosal *LCN2*-positive area (**a**) and *MARCO*-positive area (**b**) in rabbits infected with WT or Δ*zitB* at 8 hpi (*n* = 6 per group). **c**–**d** Normalized submucosal (**c**) and muscle (**d**) *LCN2*-positive area in rabbits infected with WT or Δ*zitB* at 24 hpi (*n* = 9 for WT and *n* = 8 for Δ*zitB*). **e–f** Normalized submucosal (**e**) and muscle (**f**) *MARCO*-positive area in rabbits infected with WT, Δ*zitB*, or Δ*zitB*-comp at 24 hpi (*n* = 9 for WT, *n* = 8 for Δ*zitB*, and *n* = 5 for Δ*zitB*-comp). Each point represents the model-predicted mean for one rabbit, shown on the untransformed scale. Boxes show the median and IQR; whiskers extend to the smallest and largest observations within 1.5 × IQR of the lower and upper quartiles, respectively. Data were analyzed using linear mixed-effects models with strain as a fixed effect and rabbit as a random intercept. Models for **a**–**c** were fitted to untransformed values, whereas models for **d**–**f** were fitted to log_10_-transformed values; Pairwise comparisons in **e**–**f** were Tukey-adjusted. \**P* < 0.05; ns, not significant.

**Supplementary Fig. 8:**
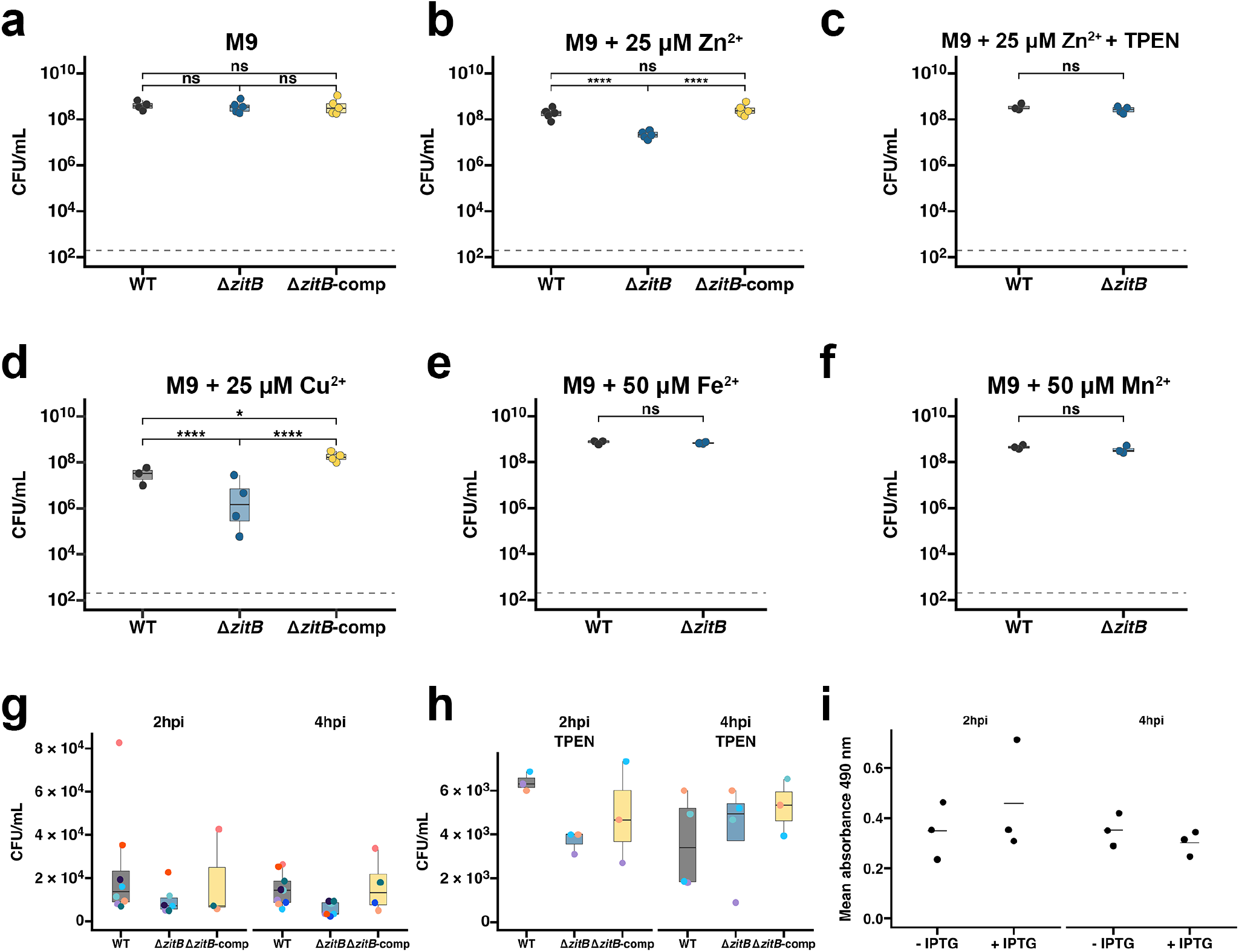
Metal-dependent growth and additional analyses of THP-1 macrophage infection. **a**–**f** Endpoint CFU after overnight growth in M9 medium without added metal (**a**) or supplemented with 25 μM Zn^2+^ (**b**), 25 μM Zn^2+^ and 25 μM TPEN (**c**), 25 μM Cu^2+^ (**d**), 50 μM Fe^2+^ (**e**), or 50 μM Mn^2+^ (**f**). Each point represents the mean CFU from one independent experiment. Sample sizes were: **a**, *n* = 4, 5, and 5 for WT, Δ*zitB*, and Δ*zitB*-comp, respectively; **b**, *n* = 5 per group; **c**, *n* = 3 and 4 for WT and Δ*zitB*; **d**, *n* = 3, 4, and 4, for WT, Δ*zitB*, and Δ*zitB*-comp respectively; and **e**–**f**, *n* = 3 per group. Dashed lines indicate the limit of detection, 2 × 10^2^ CFU ml^-1^. Data were analyzed using linear models fitted to log_10_-transformed values; pairwise comparisons in **a**, **b**, and **d** were Tukey-adjusted. **g**–**h** Raw CFU measurements from the THP-1 macrophage infection experiments shown in Fig. 5a–b, performed without TPEN (**g**) or with TPEN (**h**) at 2 and 4 hpi. Each point represents one independent infection experiment, and colors indicate experimental batch. In **g**, at 2 and 4 hpi, respectively, *n* = 8 and 9 for WT, *n* = 6 and 7 for Δ*zitB*, and *n* = 3 and 4 for Δ*zitB*-comp. In **h**, at 2 and 4 hpi, respectively, *n* = 3 and 4 for WT, 3 and 4 for Δ*zitB*, and 3 and 3 for Δ*zitB*-comp. For **a**–**h**, boxes show the median and IQR; whiskers extend to the smallest and largest observations within 1.5 × IQR of the lower and upper quartiles, respectively. **i** Mean absorbance at 490 nm for spontaneous LDH release controls consisting of uninfected THP-1 macrophages incubated with or without IPTG at 2 and 4 hpi. Each point represents one independent infection experiment (*n* = 3), and horizontal lines indicate the mean. \**P* < 0.05; \*\*\*\**P* < 0.0001; ns, not significant.

**Supplementary Fig. 9:**
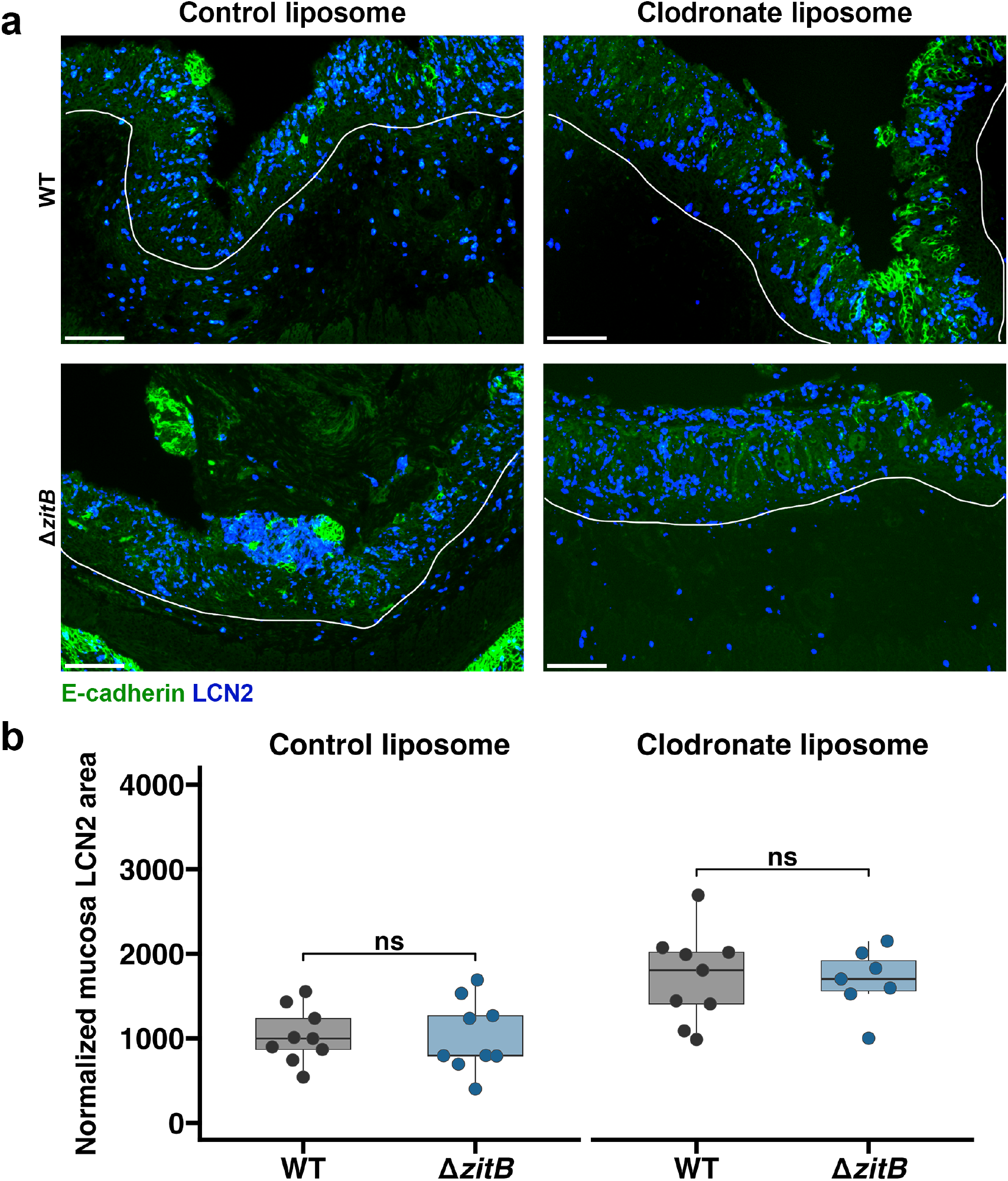
WT and Δ*zitB* infections show comparable mucosal *LCN2* signal following clodronate liposome treatment. **a** Representative images of colonic tissue from rabbits treated intravenously with control or clodronate liposomes 48 h before infection with WT or Δ*zitB* and collected at 24 hpi. *LCN2* is shown in blue and E-cadherin in green. White lines delineate the boundary between the mucosa and submucosa. Scale bars, 100 μm. **b** Quantification of normalized mucosal *LCN2*-positive area following control or clodronate liposome treatment. Each dot represents the model-predicted mean for one rabbit. Sample sizes were *n* = 9 for WT and *n* = 8 for Δ*zitB* following control liposome treatment and *n* = 9 for WT and *n* = 7 for Δ*zitB* following clodronate liposome treatment. Data were analyzed using a linear mixed-effects model strain and treatment as fixed effects and rabbit as a random intercept. The same strain effect estimated by the additive model is indicated in both treatment panels. Boxes show the median and IQR; whiskers extend to the smallest and largest observations within 1.5 × IQR of the lower and upper quartiles, respectively. ns, not significant.

## Notes

### Competing Interest Statement

The authors have declared no competing interest.

